# Firrm functions as a multiprotein interaction platform that coordinates Mcm8–9–Hrob recruitment and Fignl1-mediated Rad51 unloading

**DOI:** 10.64898/2026.10.19.760033

**Authors:** Daichi Nishiguchi, Yuto Takahashi, Naoki Tani, Yuta Nishizaki, Kensuke Tatsukawa, Yoshitaka Kawasoe, Kei-ichiro Ishiguro, Masato T. Kanemaki, Tatsuro S. Takahashi

## Abstract

Rad51 forms nucleoprotein filaments to catalyze homology search and strand exchange. The transition to recombination-associated DNA synthesis is accompanied by Rad51 unloading and recombination helicase recruitment. However, how these reactions are coordinated remains unclear. Using *Xenopus* egg extracts, we identify a mechanistic coupling between Rad51 unloading and recombination helicase recruitment. Proteomic analysis revealed that Rad51 filaments assembled on double-stranded DNA recruit a limited set of recombination regulators, including C1orf112/Firrm, Fignl1, Mcm8, Mcm9, and Hrob. Firrm forms a stoichiometric complex with Fignl1 and directly binds short acidic motifs in the intrinsically disordered regions of both Mcm9 and Hrob. The Firrm–Mcm9 interaction promotes recruitment of Mcm8–9 and Hrob to Rad51 filaments. Purified Mcm8–9 stimulates the ATPase activity of Fignl1 through Firrm in a DNA-dependent manner. Rad51 unloading from double-stranded DNA depends primarily on Fignl1 ATPase activity and is stimulated by Firrm–Mcm9 interaction. In human cells, FIRRM–FIGNL1 and MCM8–9–HROB act in the same genetic pathway to confer resistance to DNA interstrand crosslinks. These results support a model in which Rad51 filaments and Firrm–Fignl1 promote recruitment of Mcm8–9–Hrob, which, in turn, stimulates Fignl1 ATPase activity, thereby coupling recombination helicase recruitment with Rad51 unloading.

## Introduction

Rad51 is an evolutionarily conserved ATPase that plays central roles in homologous recombination (HR) and replication fork protection (Kowalczykowski 2015; Sassi et al. 2026). A fundamental property of Rad51 is its ability to assemble into nucleoprotein filaments on both single-stranded DNA (ssDNA) and double-stranded DNA (dsDNA). During somatic HR, Rad51 assembles on ssDNA generated by end resection of DNA double-strand breaks (DSBs) to form the presynaptic filament, which catalyzes homology search and strand invasion. Strand invasion generates a displacement loop (D-loop), in which nucleoprotein filaments remain associated with the heteroduplex DNA as the postsynaptic filament. Rad51 also accumulates at stalled replication forks, where it protects nascent DNA from nucleolytic degradation and promotes fork reversal (Bhat and Cortez 2018).

The postsynaptic filament stabilizes the D-loop but prevents the access of the DNA synthesis machinery to the invading 3′ terminus (Morrical 2015; Andriuskevicius et al. 2018). In yeast, the ATP-dependent DNA translocase Rad54 and the helicase Srs2 play key roles in Rad51 removal (Solinger et al. 2002; Veaute et al. 2003; Krejci et al. 2003). Rad54-mediated unloading of Rad51 from D-loops exposes the invading 3′ terminus, thereby permitting PCNA loading and recruitment of DNA polymerase δ to initiate recombination-associated DNA synthesis (Li and Heyer 2009; Li et al. 2009, 2013). Srs2 likewise alleviates Rad51-mediated inhibition of PCNA loading and DNA synthesis in multiple recombination pathways (Vasianovich et al. 2017). Although vertebrates lack a clear Srs2 homolog, the Rad54 family DNA translocases Rad54 and Rad54B have been implicated in Rad51 unloading (Mason et al. 2015). In addition, multiple helicases and DNA translocases, including BLM, FBH1, PARI, and RECQL5, have been shown to disassemble Rad51 filaments, underscoring the importance of regulating nucleofilament dynamics (Bugreev et al. 2007; Simandlova et al. 2013; Moldovan et al. 2012; Hu et al. 2007; Ito et al. 2024).

Following Rad51-mediated strand invasion, recombination-associated DNA synthesis requires coordinated remodeling of recombination intermediates, a process that involves DNA helicases (Wright et al. 2018). Mcm8–9 has emerged as an important factor in homologous recombination, particularly during interstrand crosslink (ICL) repair and meiotic recombination (Griffin and Trakselis 2019; Helderman et al. 2023). Studies in diverse experimental systems have implicated Mcm8–9 at multiple stages of HR, placing it either upstream of Rad51 filament formation, where it has been proposed to promote DNA end resection and Rad51 loading, or downstream of the Rad51 step during recombination-associated DNA synthesis (Nishimura et al. 2012; Lutzmann et al. 2012; Park et al. 2013; Lee et al. 2015; Natsume et al. 2017; McKinzey et al. 2021; Beragama Arachchi et al. 2025). Mcm8 and Mcm9 form a heterodimer that assembles into a heterohexameric AAA+ helicase related to the replicative Mcm2–7 complex (Nishimura et al. 2012; Gambus and Blow 2013; Li et al. 2021; Weng et al. 2023; McKinzey et al. 2023). The helicase is activated by Hrob (also known as Mcm8IP), which binds both Mcm8 and Mcm9 and induces the conformational changes required for helicase activation (Hustedt et al. 2019; Huang et al. 2020; Wang et al. 2020; Acharya et al. 2024; Li et al. 2026). The intrinsically disordered C-terminal region of Mcm9 directly binds Rad51 through a BRC variant motif (McKinzey et al. 2021), providing a potential physical link between the Mcm8–9–Hrob helicase and Rad51 filaments. Consistent with its proposed role in HR, disruption of Mcm8, Mcm9, or Hrob causes hypersensitivity to interstrand crosslinking agents, infertility, and genome instability, while pathogenic variants in these genes are associated with human disorders including primary ovarian insufficiency and cancer predisposition (Nishimura et al. 2012; Lutzmann et al. 2012; Hustedt et al. 2019; Huang et al. 2020; Helderman et al. 2023).

Recent studies have identified the AAA+ ATPase Fignl1 as a key Rad51 remodeler in vertebrates (Yuan and Chen 2013). Fignl1 promotes Rad51 filament disassembly on both ssDNA and dsDNA through ATP-dependent remodeling and direct interaction through the Rad51 Binding Domain (FRBD) (Matsuzaki et al. 2019; Carver et al. 2025; Kuthethur et al. 2025). In vertebrates, Fignl1 forms a stoichiometric complex with C1orf112/Firrm, a novel recombination regulator that is required for Fignl1 stability and efficient Rad51 unloading (Pinedo-Carpio et al. 2023; Stok et al. 2023; Zhang et al. 2023; Zhou et al. 2023; Tischler et al. 2024). The Firrm–Fignl1 complex is evolutionarily conserved and was originally identified in plants as FIGL1–FLIP (Fernandes et al. 2018; Edogbanya et al. 2021). Consistent with its role in regulating Rad51 dynamics, disruption of either Firrm or Fignl1 causes sensitivity to interstrand crosslinking agents, meiotic recombination defects, embryonic lethality in mice, and accumulation of unresolved recombination intermediates and ultrafine anaphase bridges (Mazouzi et al. 2023; Pinedo-Carpio et al. 2023; Stok et al. 2023; Zhang et al. 2023; Zhou et al. 2023; Tischler et al. 2024; Matsuzaki et al. 2024; Zainu et al. 2024; Ito et al. 2023). However, the molecular function of Firrm has not been clearly defined, and how Rad51 unloaders and recombination helicases are coordinated during HR remains unknown.

Here, we show that Firrm functions as a multiprotein interaction platform that coordinates Mcm8–9–Hrob recruitment and Fignl1-mediated Rad51 unloading. Using a Rad51 unloading system coupled with proteomic analysis in *Xenopus* egg extracts, we define the molecular mechanism underlying this coordination.

## Results

### Identification of proteins associated with Rad51 filaments on double-stranded DNA in *Xenopus* egg extracts

To investigate the mechanism of Rad51 unloading, we took advantage of *Xenopus* egg extracts, which efficiently recapitulate Rad51-mediated homologous recombination and fork protection (Hashimoto et al. 2010; Long et al. 2011). We immobilized a 2.9 kb plasmid DNA on Sepharose beads via a site-specific biotin modification, assembled Rad51 filaments on dsDNA, and transferred the resulting Rad51 filaments into the extracts (Fig. 1A; Supplementary Fig. 1). Nearly complete Rad51 unloading was observed at loading densities below approximately 20 molecules per plasmid and 20 ng/μL DNA (Fig. 1B,C; Supplementary Fig. 2A). Further increasing Rad51 loading progressively impaired unloading, suggesting that the unloading capacity of the extract had been exceeded. Notably, 20 ng/μL DNA roughly corresponds to the nucleoplasm-to-DNA ratio in *Xenopus* embryos at the 7th–8th cleavage divisions, and 20 molecules per 2.9-kb plasmid corresponds to approximately 2 × 10^7^ molecules per haploid *Xenopus* genome. These estimates suggest that the Rad51 unloading capacity of *Xenopus* egg extracts substantially exceeds the level required to support physiological Rad51 dynamics during early embryogenesis.

**Figure 1.**
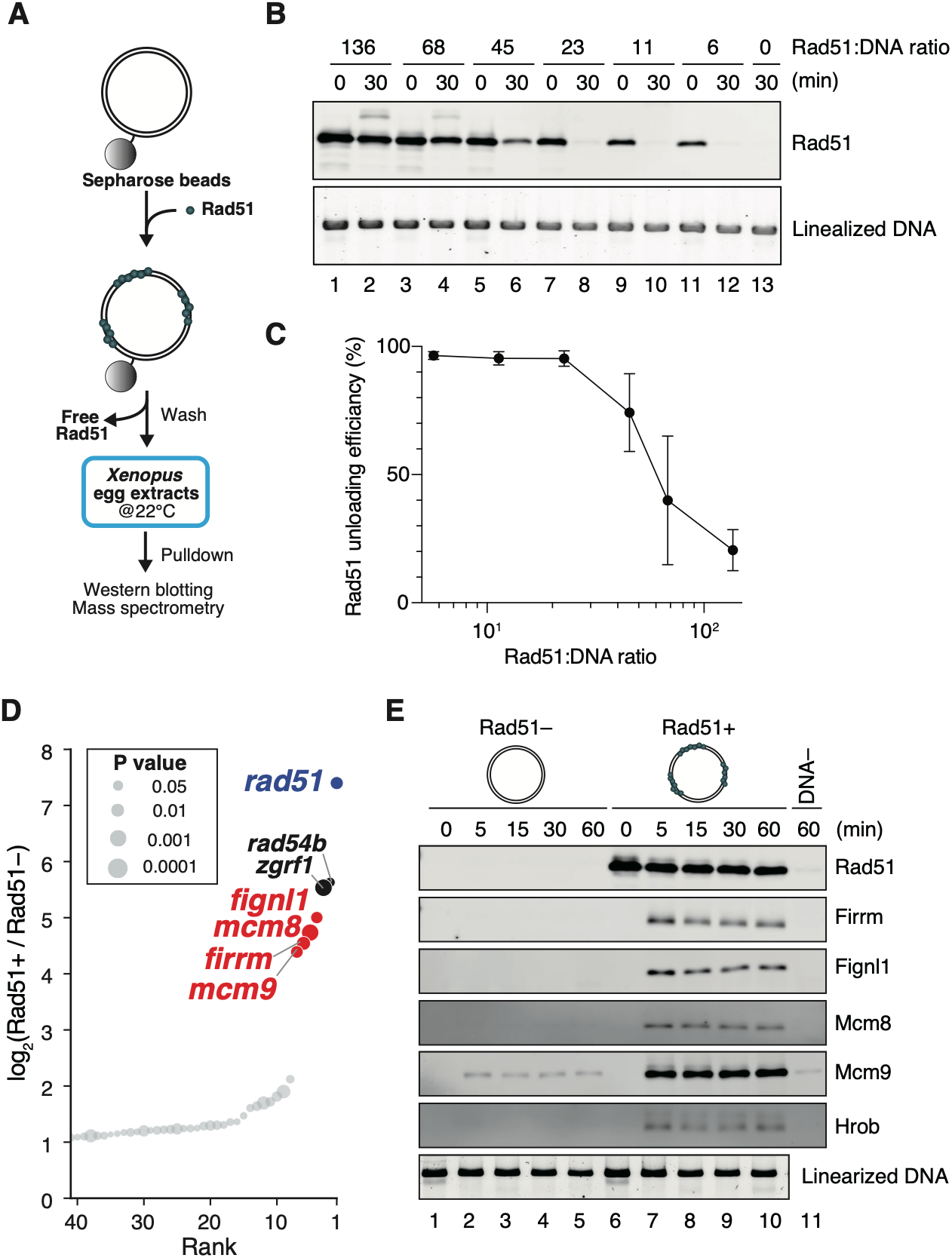
Identification of factors associated with Rad51 filaments assembled on dsDNA. (A) Schematic illustration of the Rad51 filament pull-down assay. Rad51 filaments are assembled on plasmid DNA immobilized on Sepharose beads. The Rad51–DNA complex is then incubated in *Xenopus* egg extract and recovered. (B) DNA beads carrying Rad51 filaments assembled at the indicated Rad51:DNA ratios were incubated in *Xenopus* egg extract for 30 min and recovered. Input and bead-bound fractions were analyzed by immunoblotting. Linearized DNA served as a loading control. A representative experiment from three biological replicates is shown. (C) Rad51 unloading efficiencies were quantified from replicate experiments in (B) and plotted. *n* = 3; mean ± SD. (D) Rank plot of Rad51 filament-associated factors. Rad51 filaments were assembled on dsDNA at a Rad51:DNA ratio of 68:1, and associated proteins were recovered after 30 min. Proteins are ranked according to their log₂ fold change between pull-downs with and without Rad51 filaments. Data represent three biological replicates; the top 40 proteins are shown. Circle diameter indicates the corresponding *P* value (paired *t*-test). The corresponding volcano plot is shown in Supplementary Fig. 2B. (E) Immunoblotting of Rad51 filament-associated factors. The pull-down assay was performed as described in (D). Samples were recovered at the indicated time points, separated by SDS-PAGE, and probed with the indicated antibodies. DNA− indicates the no DNA control. Linearized DNA served as a loading control. A representative experiment from three biological replicates is shown. See also Fig. 4 for related pull-down experiments that further support the reproducibility of these findings.

We next sought to identify factors responsible for Rad51 unloading. To maximize recovery of Rad51-associated proteins, we performed the analysis under conditions in which Rad51 loading exceeded the unloading capacity of the extract. Proteins whose association with dsDNA depended on Rad51 filament assembly were identified by mass spectrometry. The proteins most strongly enriched included known recombination regulators Firrm, Fignl1, Rad54B, and Zgrf1 (Fig. 1D; Supplementary Fig. 2B; Supplementary Table 1) (Wesoly et al. 2006; Yuan and Chen 2013; Brannvoll et al. 2020). Unexpectedly, the recombination helicase subunits Mcm8 and Mcm9 ranked among the most highly enriched proteins. Rad51-dependent enrichment of Firrm, Fignl1, Mcm8, and Mcm9 was confirmed by immunoblotting (Fig. 1E). Hrob, an established activator of Mcm8–9, was also enriched on Rad51-coated DNA. These findings demonstrate that Mcm8–9 and Hrob are recruited to Rad51 filaments assembled on dsDNA.

### Protein interaction network identified by pull-down analyses of Firrm–Fignl1, Mcm8–9, and Hrob

Because the DNA substrate lacks canonical helicase substrates, such as branched and forked DNA structures, recruitment of Mcm8–9 and Hrob to Rad51 filaments is likely mediated through protein–protein interactions. To investigate these interactions, we performed pull-down assays from *Xenopus* egg extracts using recombinant proteins as bait (Fig. 2A; Supplementary Fig. 1). Strikingly, Firrm and Fignl1 were among the highest-confidence hits in the Mcm8–9 pull-down (Fig. 2B; Supplementary Fig. 3A; Supplementary Table 2). Reciprocally, recombinant Firrm–Fignl1 recovered Mcm8–9 (Fig. 2C; Supplementary Fig. 3B; Supplementary Table 3). Recombinant Hrob likewise recovered both Mcm8–9 and Firrm–Fignl1, whereas Firrm–Fignl1 also recovered Hrob (Fig. 2D; Supplementary Fig. 3C; Supplementary Table 4). Immunoprecipitation from *Xenopus* egg extracts confirmed the association of endogenous Mcm8–9, Hrob, Firrm–Fignl1, and Rad51 (Fig. 2E–H). Notably, immunoprecipitation of Firrm and Fignl1 resulted in nearly stoichiometric co-recovery of the reciprocal subunit, as did immunoprecipitation of Mcm8 and Mcm9, indicating that Firrm–Fignl1 and Mcm8–9 exist predominantly as stable complexes in the extracts (Fig. 2E, lanes 2–5 and 9–12). In contrast, co-immunoprecipitation between Firrm–Fignl1, Mcm8–9, and Hrob was substoichiometric, suggesting that these factors do not exist as a stable higher-order complex. However, all three factors sedimented substantially faster than expected from their predicted molecular masses in glycerol gradients, leaving open the possibility that they form larger, dynamic assemblies in *Xenopus* egg extracts (Supplementary Fig. 3D).

**Figure 2.**
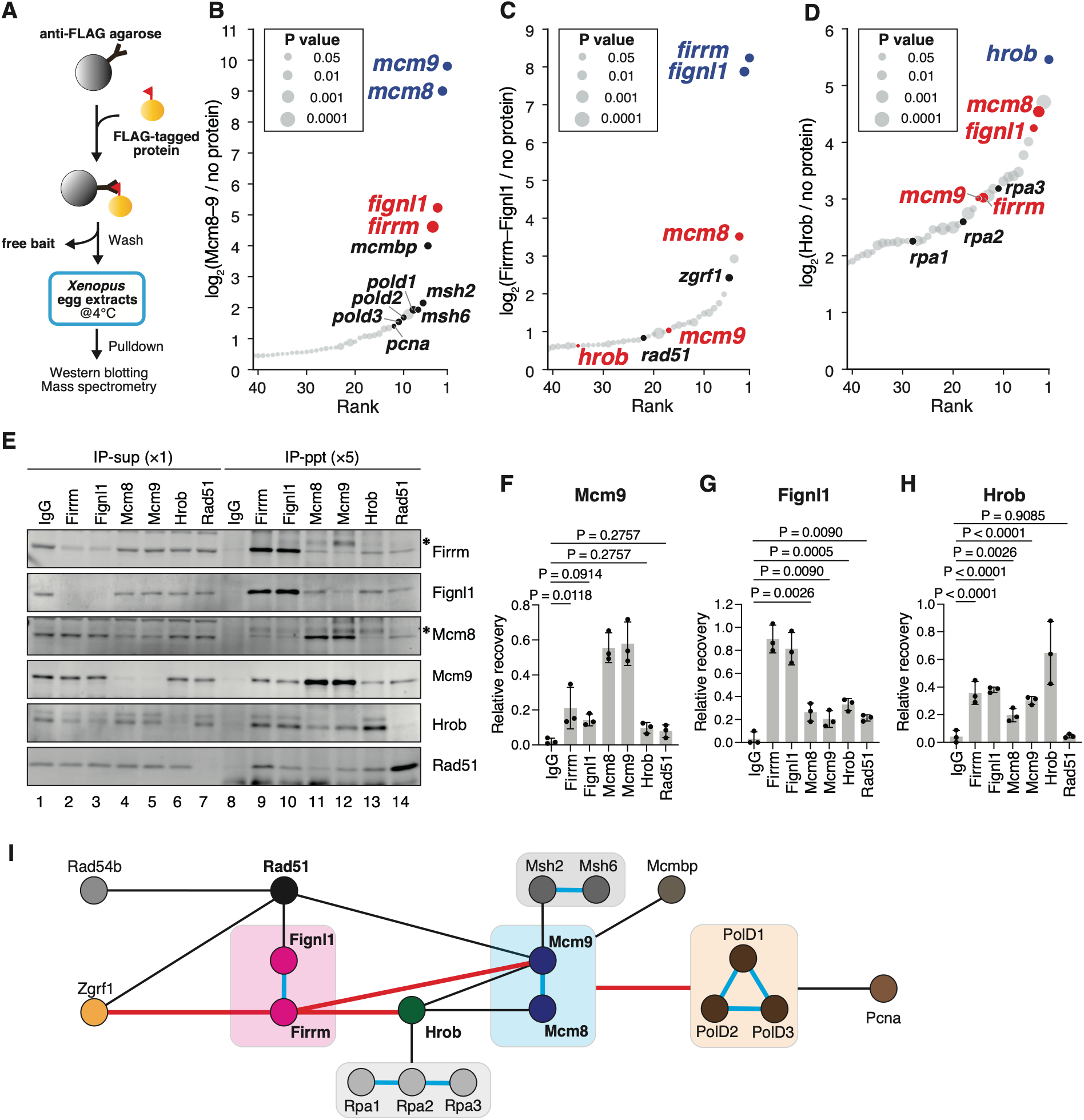
Protein interaction network identified by pull-down analyses of Firrm–Fignl1, Mcm8–9, and Hrob. (A) Schematic illustration of the recombinant protein pull-down assay. FLAG-tagged recombinant proteins are immobilized on anti-FLAG M2 agarose beads, incubated in *Xenopus* egg extract, and recovered. (B–D) Rank plots of proteins identified by pull-down assay using recombinant Mcm8–9 (B), Firrm–Fignl1 (C), and Hrob (D) as bait. Proteins are ranked according to their log₂ fold change between pull-downs with and without the corresponding bait. Data represent three biological replicates; the top 40 proteins are shown. Circle diameter indicates the corresponding *P* value (paired *t*-test). The corresponding volcano plots are shown in Supplementary Fig. 3A–C. (E) Co-immunoprecipitation analysis in *Xenopus* egg extracts. Immunoprecipitation was performed using the antibodies indicated at the top. Supernatant and immunoprecipitate fractions were loaded in the indicated proportions, separated by SDS-PAGE, and probed with the antibodies indicated on the right. A representative experiment from three biological replicates is shown. Asterisks indicate cross-reacting bands. (F–H) Quantification of immunoblot signals. Relative recovery of Mcm9 (F), Fignl1 (G), and Hrob (H) was calculated from replicate experiments in (E), with the signal in lane 1 set to 1. *n* = 3; mean ± SD; one-way ANOVA with Holm–Šídák’s multiple comparisons test. (I) Summary of the protein interaction network identified in this study. Shaded boxes, stable protein complexes; black lines, previously reported interactions; cyan lines, constitutive interactions within protein complexes; red lines, interactions identified in this study.

The mass spectrometry analyses also revealed broader interaction profiles of these proteins (Fig. 2I). Consistent with previous reports, the Mcm8–9 interactome included Msh2–Msh6 and Mcmbp (Taipale et al. 2014; Traver et al. 2015; Terui et al. 2018; Goellner et al. 2018). The Mcm8–9 pull-down also recovered multiple Polδ subunits and PCNA, suggesting a functional connection between this recombination helicase and the DNA synthesis machinery. The Hrob pull-down identified Rpa1–3, as reported previously (Huang et al. 2020). In addition to Mcm8–9 and Hrob, the Firrm–Fignl1 pull-down was enriched for the Zgrf1 helicase together with Rad51, suggesting a possible functional relationship between Zgrf1 and Firrm–Fignl1. The distinct interaction profiles of Firrm–Fignl1, Mcm8–9, and Hrob further support the conclusion that these proteins are not constitutively assembled into a higher-order complex, despite their extensive physical interactions.

### Mcm8–9 and Hrob directly interact with Firrm via short motifs on their IDRs

To gain insights into the interaction between Mcm8–9, Hrob, and Firrm–Fignl1, we used AlphaFold3 (AF3) to predict their interaction interfaces (Abramson et al. 2024). Consistent with a previous AlphaFold2-Multimer model (Stok et al. 2023), AF3 predicted that Firrm and Fignl1(1–143) form a compact, predominantly α-helical complex, in which three α-helices from Fignl1 are integrated into the structural core of Firrm (Fig. 3A). AF3 further predicted that both Mcm9 and Hrob engage a common positively charged surface cleft of Firrm through short acidic motifs located within their intrinsically disordered regions (IDRs) (Fig. 3A,B; Supplementary Fig. 4A,B). The predicted Mcm9-binding motif resides within the C-terminal IDR (residues 1116–1132), whereas the corresponding Hrob motif resides within the N-terminal IDR (residues 2–31). In both cases, evolutionarily conserved hydrophobic residues (Supplementary Fig. 4C,D) are predicted to dock into hydrophobic pockets within the Firrm surface cleft, likely stabilizing the interaction. An AlphaFold-Multimer-based in silico screen independently identified both the human MCM9–FIRRM and HROB–FIRRM interactions, providing additional support for the conservation of these interactions across species (Schmid and Walter 2025). Interestingly, unlike Mcm9 and Hrob, Zgrf1 is predicted to bind a distinct acidic surface of Firrm through a conserved basic peptide in its N-terminal IDR (Supplementary Fig. 5). These predicted interactions suggest that Firrm functions as a molecular interaction platform for multiple recombination regulators.

**Figure 3.**
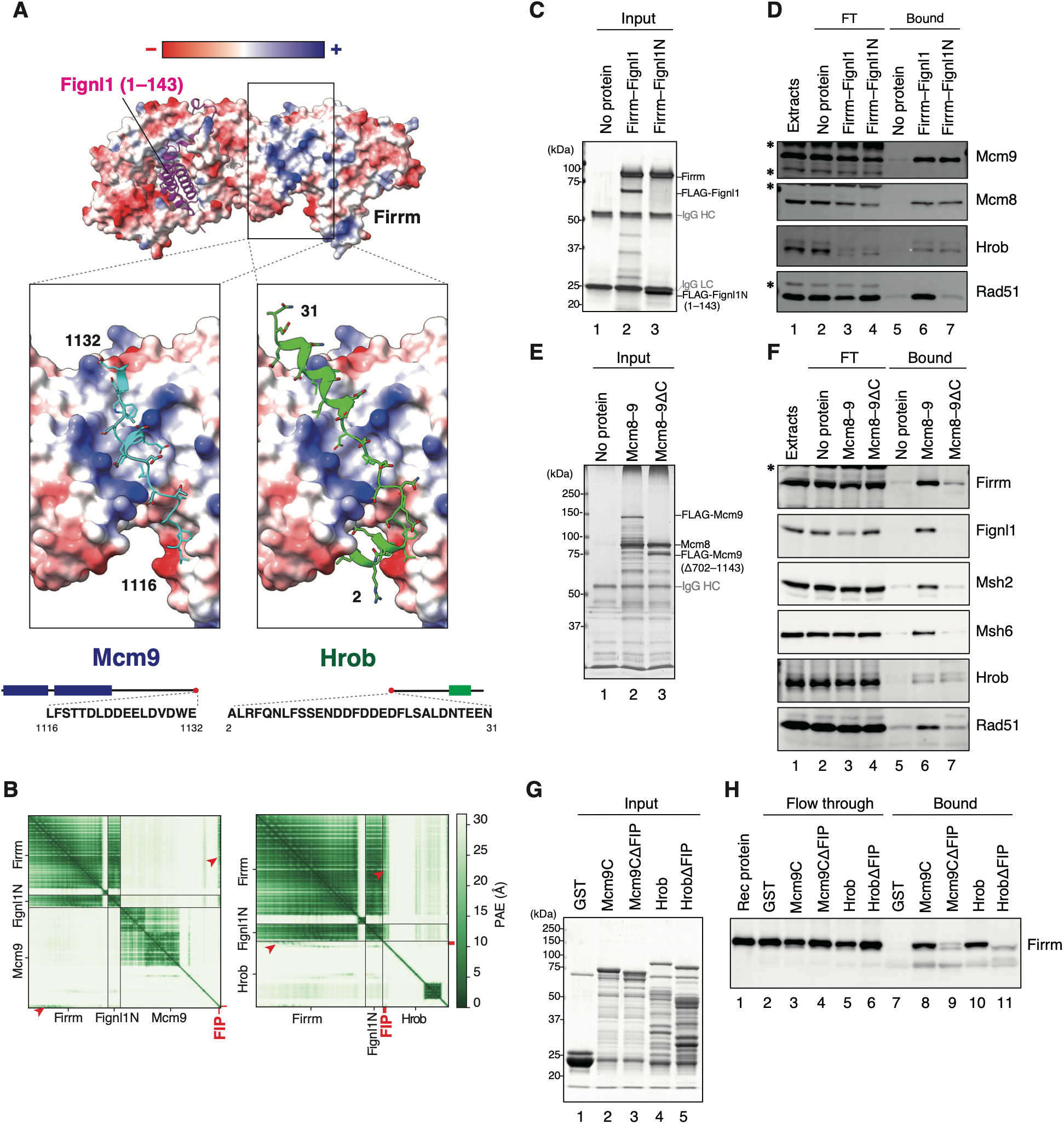
Mcm8–9 and Hrob directly interact with Firrm via short motifs on their IDRs. (A) AlphaFold3-predicted structures of the Firrm–Fignl1(1–143) complex bound to Mcm9- or Hrob-derived peptides. The surface of Firrm is colored according to its electrostatic surface potential, with basic and acidic regions shown in blue and red, respectively. Mcm9- and Hrob-derived peptides are shown in cyan and green, respectively. Residue ranges and amino acid sequences of the peptides are shown below. (B) Predicted aligned error (PAE) heat maps for the AlphaFold3 models shown in (A). Red arrowheads indicate the predicted Firrm–Mcm9 and Firrm–Hrob interfaces. (C, D) Pull-down assay using recombinant Firrm–Fignl1 and Firrm–Fignl1N complexes. Recombinant proteins were immobilized on anti-FLAG M2 agarose beads and used as bait for pull-down from *Xenopus* egg extracts. Silver staining of input samples (C) and immunoblot analysis of flow-through (FT) and bound fractions probed with the indicated antibodies (D) are shown. A representative experiment from three biological replicates is shown. IgG HC, immunoglobulin heavy chain; IgG LC, immunoglobulin light chain. Asterisks indicate cross-reacting bands. Statistical analyses are shown in Supplementary Fig. 6A–C. (E, F) Pull-down assay using recombinant Mcm8–9 and Mcm8–Mcm9ΔC (Δ702–1143). The assay was performed as described in (C, D). Silver staining of input samples (E) and immunoblot analysis of flow-through (FT) and bound fractions (F) are shown. A representative experiment from three biological replicates is shown. Statistical analyses are shown in Supplementary Fig. 6D–G. (G, H) Recombinant pull-down assay using GST, GST-Mcm9C [Mcm9(702–1143)], GST-Mcm9CΔFIP [Mcm9(702–1143, Δ1116–1132)], GST-Hrob, and GST-HrobΔFIP [Hrob(Δ2–31)]. GST fusion proteins were immobilized on Glutathione Sepharose beads, incubated with recombinant Firrm–Fignl1N, and recovered. Coomassie Brilliant Blue R-250 staining of input samples (G) and immunoblot analysis of flow-through and bound fractions probed with Firrm antibodies (H) are shown. A representative experiment from three technical replicates is shown. Statistical analyses are shown in Supplementary Fig. 6H.

To test whether Firrm–Fignl1 directly interacts with the predicted Mcm9 and Hrob motifs, we performed in vitro pull-down assays. Consistent with the AF3 model, Firrm was insoluble when expressed alone but became soluble upon co-expression with Fignl1(1–143), allowing purification of the resulting Firrm–Fignl1(1–143) complex, hereafter referred to as Firrm–Fignl1N (Supplementary Fig. 1). Before using this complex in reconstituted binding assays, we confirmed by pull-down assays in *Xenopus* egg extracts that Firrm–Fignl1N retained the ability to interact with both Mcm8–9 and Hrob (Fig. 3C,D; Supplementary Fig. 6A–C). Using Flag-tagged Mcm8–9 as bait, we also found that the C-terminal IDR of Mcm9 (Mcm9C) is required for the interaction with Firrm–Fignl1 (Fig. 3E,F; Supplementary Fig. 6D). Mcm9C contains the Msh2 interaction motif (Goellner et al. 2018) and a BRC variant motif that binds Rad51 (McKinzey et al. 2021), and we confirmed that Mcm9C is required for the association with both Msh2–Msh6 and Rad51, but not with Hrob (Fig. 3F; Supplementary Fig. 6E–G). The Mcm9C fragment, full-length Hrob, and shorter peptides containing the predicted Firrm-interacting motifs were then expressed in bacteria as GST fusion proteins and used for GST pull-down assays with purified Firrm–Fignl1N (Fig. 3G; Supplementary Fig. 6I). Consistent with the AF3 predictions, both Mcm9 and Hrob directly bound Firrm–Fignl1N, whereas deletion of the respective predicted interaction motifs largely abolished binding (Fig. 3H; Supplementary Fig. 6H, J, K). These results validate the AF3-predicted Firrm-binding motifs in Mcm9 and Hrob. We hereafter refer to these motifs as Firrm-interacting peptide (FIP) motifs.

### Firrm–Fignl1 promotes recruitment of Mcm8–9–Hrob to Rad51 filaments

We next investigated how these interactions contribute to the recruitment of each factor to Rad51 filaments. Depletion of Fignl1 from the extracts reduced the accumulation of both Mcm8–9 and Hrob on Rad51 filaments by approximately one-half to two-thirds (Fig. 4A,B; Supplementary Fig. 7A–C). Rescue experiments with recombinant Firrm–Fignl1 showed that accumulation of Firrm on Rad51 filaments requires the 144–655 fragment of Fignl1 but not its ATPase motif, as the Walker A-mutant Firrm–Fignl1(K428A), but not Firrm–Fignl1N, restored Firrm accumulation (Fig. 4C,D; Supplementary Fig. 7D). Likewise, both wild-type and Walker A-mutant Firrm–Fignl1, but not Firrm–Fignl1N, stimulated accumulation of Mcm8, Mcm9, and Hrob, indicating that the Firrm–Fignl1-dependent fraction of Mcm8–9 and Hrob recruitment requires the Fignl1 fragment but not its ATPase motif (Fig. 4C–F; Supplementary Fig. 7E).

**Figure 4.**
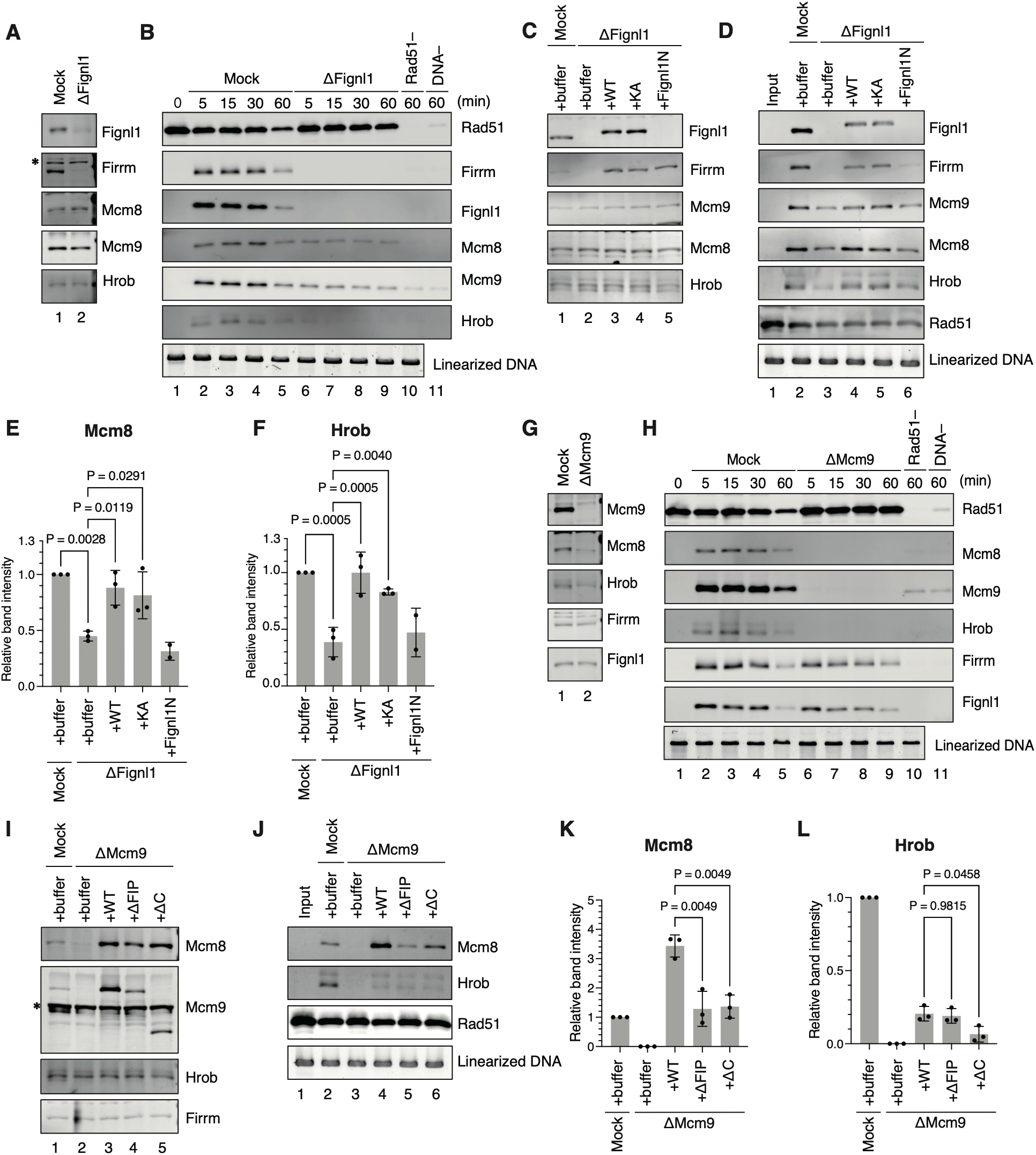
Firrm–Fignl1 promotes Mcm8–9–Hrob recruitment to Rad51 filaments through the Firrm–Mcm9 interaction. (A) Immunoblot analysis of mock- and Fignl1-depleted extracts. The extracts were separated by SDS-PAGE and probed with the indicated antibodies. Asterisk indicates cross-reacting band. (B) Rad51 filaments were assembled on immobilized DNA at a Rad51:DNA ratio of 68:1, incubated in the extracts shown in (A), and recovered at the indicated time points. Bead-bound proteins were separated by SDS-PAGE and probed with the indicated antibodies. Linearized DNA served as a loading control. A representative experiment from three biological replicates is shown. Statistical analyses are shown in Supplementary Fig. 7A–C. (C) Immunoblot analysis of Fignl1 depletion/add-back experiments. Recombinant proteins were added back to a final concentration of 90 nM. Buffer, control buffer; WT, wild-type Firrm–Fignl1; KA, Firrm–Fignl1(K428A); Fignl1N, Firrm–Fignl1(1–143). (D) Rad51 filament pull-down assay using the extracts from (C). Rad51 filaments were assembled on dsDNA at a Rad51:DNA ratio of 68:1. Samples were recovered at 30 min, separated by SDS-PAGE, and probed with the indicated antibodies. A representative experiment from three biological replicates is shown. (E, F) Quantification of immunoblot signals. Relative band intensities of Mcm8 (E) and Hrob (F) were calculated from replicate experiments in (D), with the signal in lane 2 set to 1. *n* = 3; mean ± SD; one-way ANOVA with Šídák’s multiple comparisons test. Statistical analyses of Firrm and Mcm9 are shown in Supplementary Fig. 7D and E. (G) Immunoblot analysis of mock- and Mcm9-depleted extracts. (H) Rad51 filament pull-down assay using the extracts from (G). The pull-down assay and immunoblot analysis were performed as described in (B). A representative experiment from three biological replicates is shown. Statistical analyses are shown in Supplementary Fig. 7F and G. (I) Immunoblot analysis of Mcm9 depletion/add-back experiments. Recombinant Mcm8–9 proteins were added back to a final concentration of 160 nM. Buffer, control buffer; WT, wild-type Mcm8–9; ΔFIP, Mcm8–Mcm9ΔFIP [Mcm9(Δ1116–1132)]; ΔC, Mcm8–Mcm9ΔC [Mcm9(Δ702–1143)]. Asterisk indicates cross-reacting band. (J) Rad51 filament pull-down assay using the extracts from (I). The pull-down assay and immunoblot analysis were performed as described in (D). A representative experiment from three biological replicates is shown. (K, L) Quantification of immunoblot signals. The relative band intensity of Mcm8 (K) and Hrob (L) was calculated from replicate experiments in (J), with the signal in lane 2 set to 1. *n* = 3; mean ± SD; one-way ANOVA with Šídák’s multiple comparisons test.

In contrast, depletion of Mcm8–9 did not significantly affect the accumulation of Firrm–Fignl1 (Fig. 4G,H; Supplementary Fig. 7F,G). Depletion of Mcm9, which partially co-depleted Hrob, almost completely inhibited Hrob accumulation on Rad51 filaments (Fig. 4G,H). Rescue experiments with recombinant Mcm8–9 (Supplementary Fig. 1) showed that deletion of either the FIP motif or the C-terminal IDR of Mcm9 reduced Mcm8–9 loading by approximately one-half to two-thirds, strongly suggesting that Firrm recruits Mcm8–9 to Rad51 filaments through the FIP-mediated interaction (Fig. 4I–K). Hrob recruitment was only partially restored, likely because Hrob was partially co-depleted with Mcm9 (Fig. 4L). Collectively, these findings define a recruitment hierarchy in which recruitment of Firrm requires Fignl1, Firrm–Fignl1 mediates recruitment of a substantial fraction of Mcm8–9, and Mcm8–9 in turn stabilizes Hrob recruitment to Rad51 filaments.

### Interaction with Mcm8–9 stimulates the Fignl1 ATPase in a DNA-dependent manner

Since Fignl1, Mcm8, and Mcm9 are ATPases, we next asked whether their physical interactions influence ATP hydrolysis by these proteins. To address this, we performed in vitro ATPase assays in the presence or absence of a 70-mer dA/dT oligonucleotide. Both Mcm8–9 and Firrm–Fignl1 exhibited basal levels of ATP hydrolysis in the absence of DNA, whereas DNA did not strongly stimulate their ATPase activities under our assay conditions (Fig. 5A, compare columns 3 and 4, and columns 7 and 8, respectively). In contrast, co-incubation of Mcm8–9 and Firrm–Fignl1 resulted in robust, DNA-dependent stimulation of ATPase activity (Fig. 5A, columns 11 and 12). Deletion of the C-terminal tail or the FIP motif in Mcm9 abolished ATPase stimulation, demonstrating that stimulation requires the interaction between Mcm8–9 and Firrm–Fignl1 (Fig. 5B, compare columns 5 and 8, and Supplementary Fig. 8A). Although Hrob interacts with both complexes, its addition did not further enhance ATP hydrolysis by either complex or by the combined reaction (Fig. 5A, columns 10, 14 and 16). These findings suggest that complex formation between Mcm8–9 and Firrm–Fignl1 stimulates ATP hydrolysis by Mcm8–9, Firrm–Fignl1, or both.

**Figure 5.**
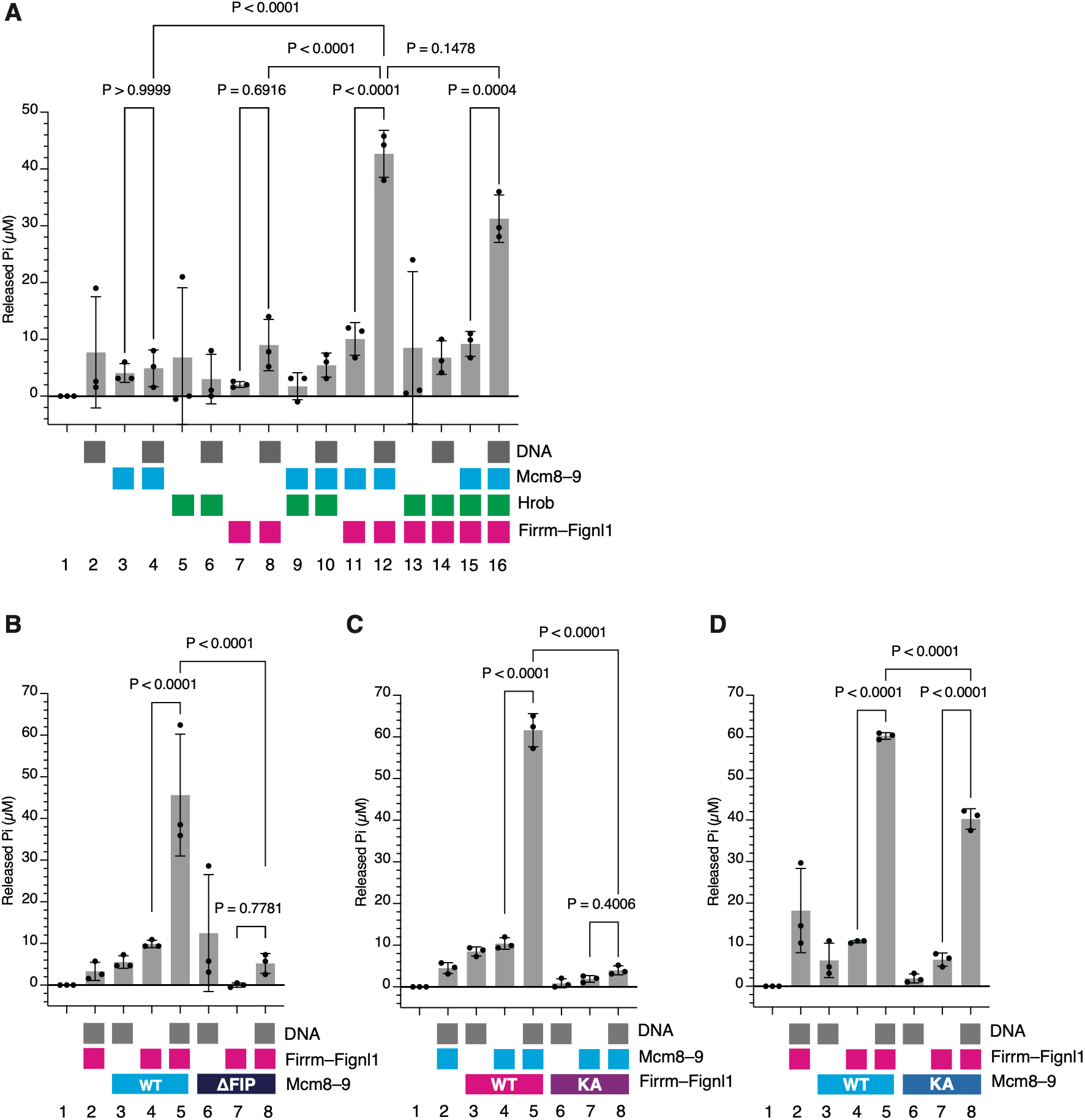
Mcm8–9 activates the Fignl1 ATPase in a manner dependent on both DNA and the Firrm–Mcm9 interaction. (A–D) ATPase activity was quantified by measuring released inorganic phosphate. Recombinant proteins were used at 50 nM each: Mcm8–9, Hrob, and Firrm–Fignl1 in (A); Firrm–Fignl1, Mcm8–9, and Mcm8–9ΔFIP [Mcm9(Δ1116–1132)] in (B); Mcm8–9, Firrm–Fignl1, and Firrm–Fignl1(K428A) in (C); and Firrm–Fignl1, Mcm8–9, and Mcm8(K453A)–Mcm9(K359A) in (D). Poly-dA/dT DNA was added at 1 μM. Squares indicate the proteins and DNA present in each reaction. *n* = 3; mean ± SD; one-way ANOVA with Šídák’s multiple comparisons test.

To determine which ATPase is stimulated, we mutated the Walker A motifs of Mcm8(K453A), and Mcm9(K359A), as well as Fignl1(K428A) (Supplementary Fig. 1). Strikingly, the Fignl1 ATPase mutant failed to support the stimulation of ATP hydrolysis (Fig. 5C). Likewise, Firrm–Fignl1N failed to stimulate ATP hydrolysis (Supplementary Fig. 8B). In contrast, the Mcm8–9 mutant carrying Walker A substitutions in both subunits retained the ability to stimulate ATP hydrolysis when combined with wild-type Firrm–Fignl1 (Fig. 5D). These results demonstrate that interaction with Mcm8–9 stimulates the ATPase of Fignl1 in a DNA-dependent manner.

The additional release of 30–50 μM inorganic phosphate corresponds to an apparent ATP turnover of approximately 5–8 ATP molecules per minute per Fignl1 molecule. This value is likely an underestimate because ATP hydrolysis was averaged over the total Fignl1 population, whereas only a subset of molecules may have been stimulated by Mcm8–9.

### Mcm8–9 stimulates Rad51 unloading in *Xenopus* egg extracts

The above findings support a model in which Firrm–Fignl1 promotes the recruitment of Mcm8–9 and Hrob to Rad51 filaments, whereas Mcm8–9, in turn, stimulates the Fignl1 ATPase, possibly to enhance Rad51 unloading. To test this model directly, we performed a Rad51-unloading assay in *Xenopus* egg extracts using a limited amount of Rad51 filaments assembled on dsDNA (Fig.1A). We first examined whether Firrm–Fignl1 is required for Rad51 unloading. In mock-treated extracts, Rad51 was rapidly unloaded from DNA within 10 min (Fig. 6A–C). In contrast, Fignl1 depletion largely prevented Rad51 unloading. Rescue experiments with recombinant Firrm–Fignl1 showed that Rad51 unloading requires an intact ATPase motif in Fignl1 (Fig. 6D–F). Consistent with a previous report (Carver et al. 2025), these results demonstrate that the ATPase activity of Fignl1 is crucial for Rad51 unloading from dsDNA in *Xenopus* egg extracts.

**Figure 6.**
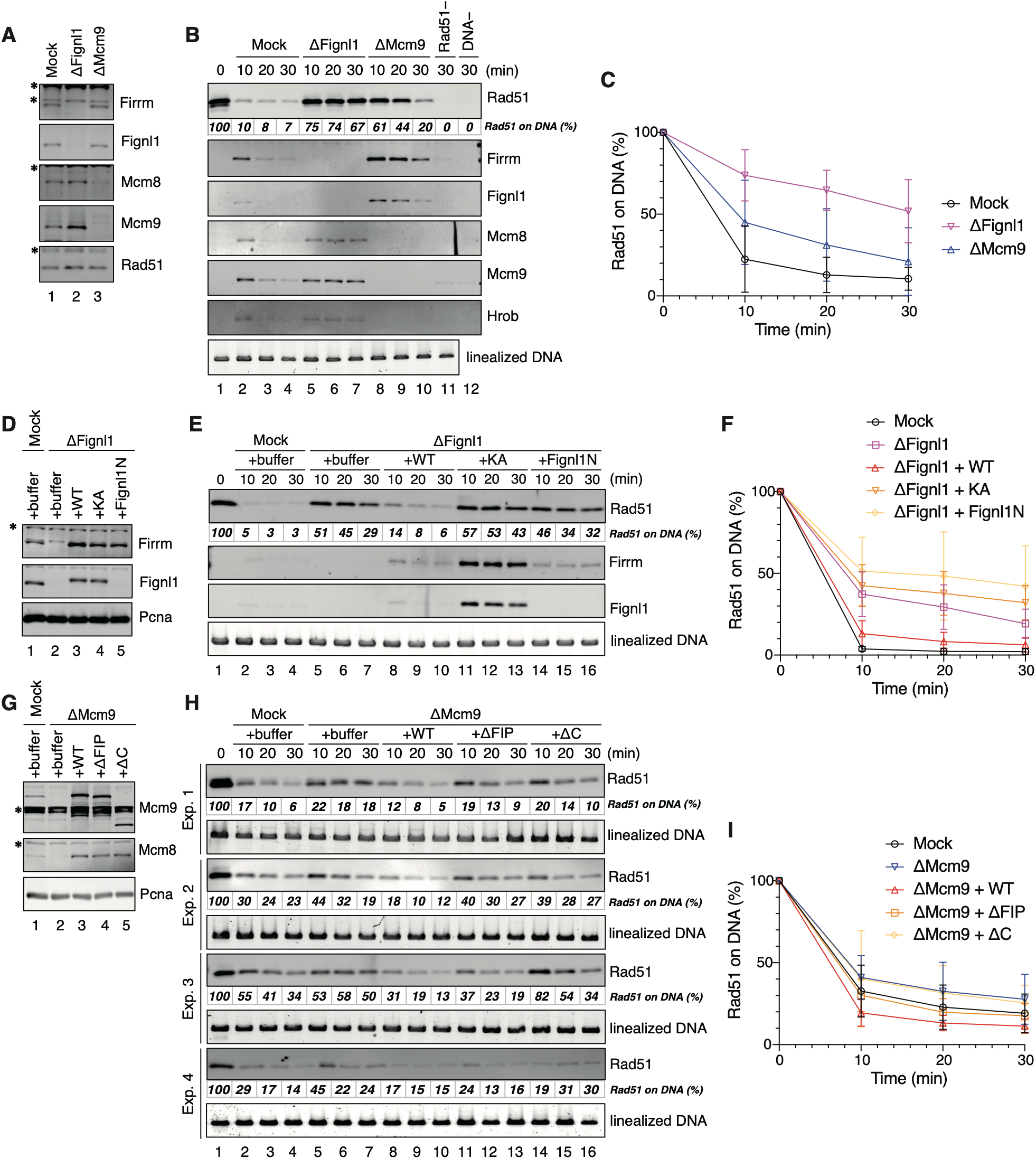
Mcm8–9 promotes Rad51 unloading through the Mcm9–Firrm interaction in *Xenopus* egg extracts. (A) Immunoblot analysis of mock, Fignl1-depleted, and Mcm9-depleted extracts. Asterisks indicate cross-reacting bands. (B) Rad51 filament pull-down assay using the extracts from (A). Rad51 filaments were assembled on dsDNA at a Rad51:DNA ratio of 23:1. Samples were recovered at the indicated time points, separated by SDS-PAGE, and probed with the indicated antibodies. Values below the Rad51 blot indicate the percentage of Rad51 remaining, with lane 1 set to 100%. Linearized DNA served as a loading control. A representative experiment from four biological replicates is shown. (C) Quantification of Rad51 signals from replicate experiments in (B). *n* = 4; mean ± SD. (D) Immunoblot analysis of Fignl1 depletion/add-back experiments. Recombinant proteins were added back to a final concentration of 90 nM. Buffer, control buffer; WT, wild-type Firrm–Fignl1; KA, Firrm–Fignl1(K428A); Fignl1N, Firrm–Fignl1(1–143). Asterisk indicates cross-reacting band. (E) Rad51 filament pull-down assay using the extracts from (D). Rad51 filaments were assembled on dsDNA at a Rad51:DNA ratio of 3:1. A representative experiment from three biological replicates is shown. (F) Quantification of Rad51 signals from replicate experiments in (E). *n* = 3; mean ± SD. (G) Immunoblot analysis of Mcm9 depletion/add-back experiments. Recombinant Mcm8–9 proteins were added back to a final concentration of 160 nM. Buffer, control buffer; WT, wild-type Mcm8–9; ΔFIP, Mcm8–Mcm9ΔFIP [Mcm9(Δ1116–1132)]; ΔC, Mcm8–Mcm9ΔC [Mcm9(Δ702–1143)]. Asterisks indicate cross-reacting bands. (H) Rad51 filament pull-down assay using the extracts from (G). Rad51 filaments were assembled on dsDNA at a Rad51:DNA ratio of 11:1. Four biological replicates (Exp. 1–4) are shown. (I) Quantification of Rad51 signals from replicate experiments in (H). *n* = 4; mean ± SD.

Strikingly, we found that Mcm9 depletion also reduced the rate of Rad51 unloading (Fig. 6A–C). One-phase exponential decay fitting estimated that Mcm9 depletion extended the half-life of Rad51 retention approximately twofold, whereas Fignl1 depletion extended it approximately fivefold (Table 1,2). To test whether Mcm8–9 stimulates Rad51 unloading through interaction with Firrm, we immunodepleted Mcm9 from the extracts and supplemented them with either wild-type Mcm8–9 or mutants lacking the FIP motif or the C-terminal IDR (Fig. 6G). Because Mcm9 depletion increases the estimated half-life of Rad51 unloading by only approximately twofold, the dynamic range of the complementation assay is inherently limited. Nevertheless, wild-type Mcm8–9 stimulated Rad51 unloading, whereas neither the ΔFIP nor the ΔC mutant restored unloading to the level observed with wild-type Mcm8–9 in any of four independent experiments (Fig. 6H,I; Table 3). Together with the biochemical interaction and ATPase data presented above, these findings support a model in which the interaction between Mcm8–9 and Firrm stimulates Rad51 unloading.

**Table 1.** Fitting parameters for the Rad51 unloading assay.

| Sample | $t_{1/2}$ (min) | [95% Confidence Interval] | $R^2$ |
| --- | --- | --- | --- |
| mock | 5.4 | [3.5–7.3] | 0.91 |
| ΔFigl1 | 30 | [23–41] | 0.66 |
| ΔMcm9 | 11 | [7.4–16] | 0.73 |

**Table 2.**
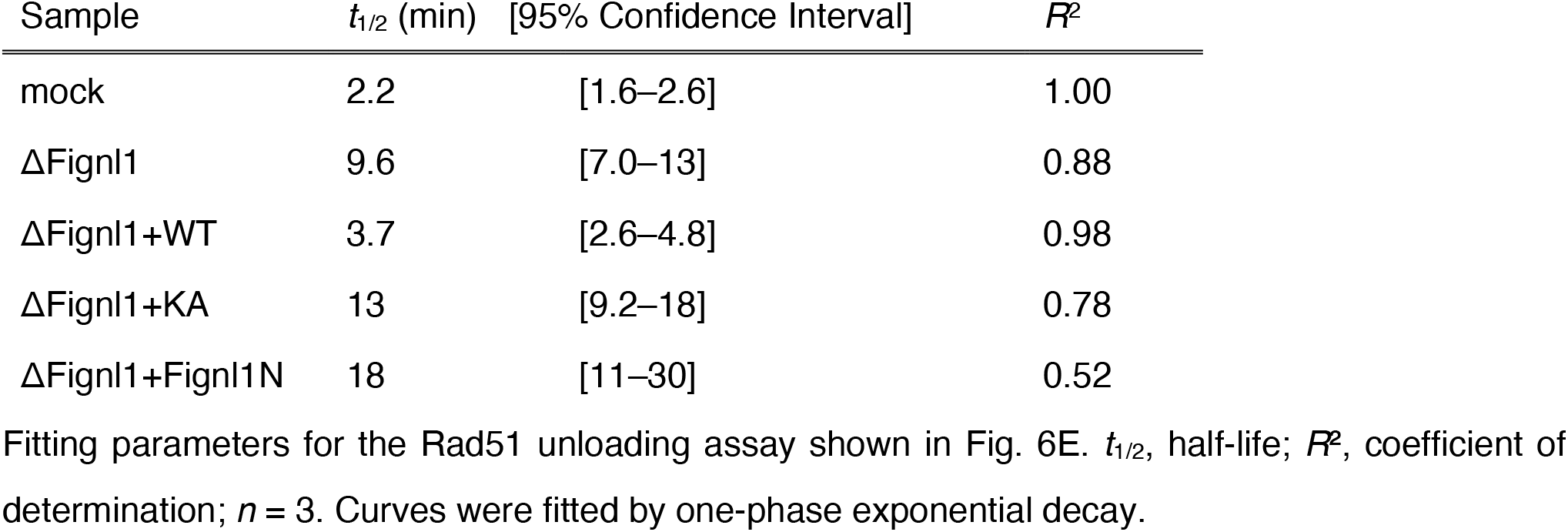
Fitting parameters for the Rad51 unloading assay following Firrm–Fignl1 depletion and add-back.

| Sample | $t_{1/2}$ (min) | [95% Confidence Interval] | $R^2$ |
| --- | --- | --- | --- |
| mock | 2.2 | [1.6–2.6] | 1.00 |
| ΔFigl1 | 9.6 | [7.0–13] | 0.88 |
| ΔFigl1+WT | 3.7 | [2.6–4.8] | 0.98 |
| ΔFigl1+KA | 13 | [9.2–18] | 0.78 |
| ΔFigl1+Figl1N | 18 | [11–30] | 0.52 |

**Table 3.** Fitting parameters for the Rad51 unloading assay following Mcm8–9 depletion and add-back.

| Sample | $t_{1/2}$ (min) | [95% Confidence Interval] | $R^2$ |
| --- | --- | --- | --- |
| mock | 8.2 | [6.2–10] | 0.86 |
| ΔMcm9 | 11 | [8.8–15] | 0.78 |
| ΔMcm9+WT | 5.0 | [3.9–6.0] | 0.95 |
| ΔMcm9+ΔFIP | 7.4 | [5.9–9.0] | 0.92 |
| ΔMcm9+ΔC | 11 | [7.9–15] | 0.72 |

### Firrm–Fignl1 and Mcm8–9–Hrob function in the same pathway for resistance to DNA interstrand crosslinks in human cells

To investigate the relationship between Firrm–Fignl1 and Mcm8–9–Hrob in human cells, we knocked out these genes in HCT116 colorectal cancer cells using CRISPR-Cas9 (see Materials & Methods) and assessed their sensitivity to the DNA interstrand crosslinking agent mitomycin C (MMC). Consistent with previous reports, *FIGNL1*, *FIRRM*, *MCM8* and *HROB* knockout (KO) cells were more sensitive to MMC than HCT116 WT cells (Fig. 7A–C) (Nishimura et al. 2012; Hustedt et al. 2019; Huang et al. 2020; Wang et al. 2020; Zhou et al. 2023; Mazouzi et al. 2023; Stok et al. 2023; Pinedo-Carpio et al. 2023; Tischler et al. 2024). However, *MCM8* and *HROB* KO cells exhibited more pronounced MMC sensitivity than *FIRRM* and *FIGNL1* KO cells. This difference in MMC sensitivity was also observed in independent KO clones (Supplementary Fig. 9).

**Figure 7.**
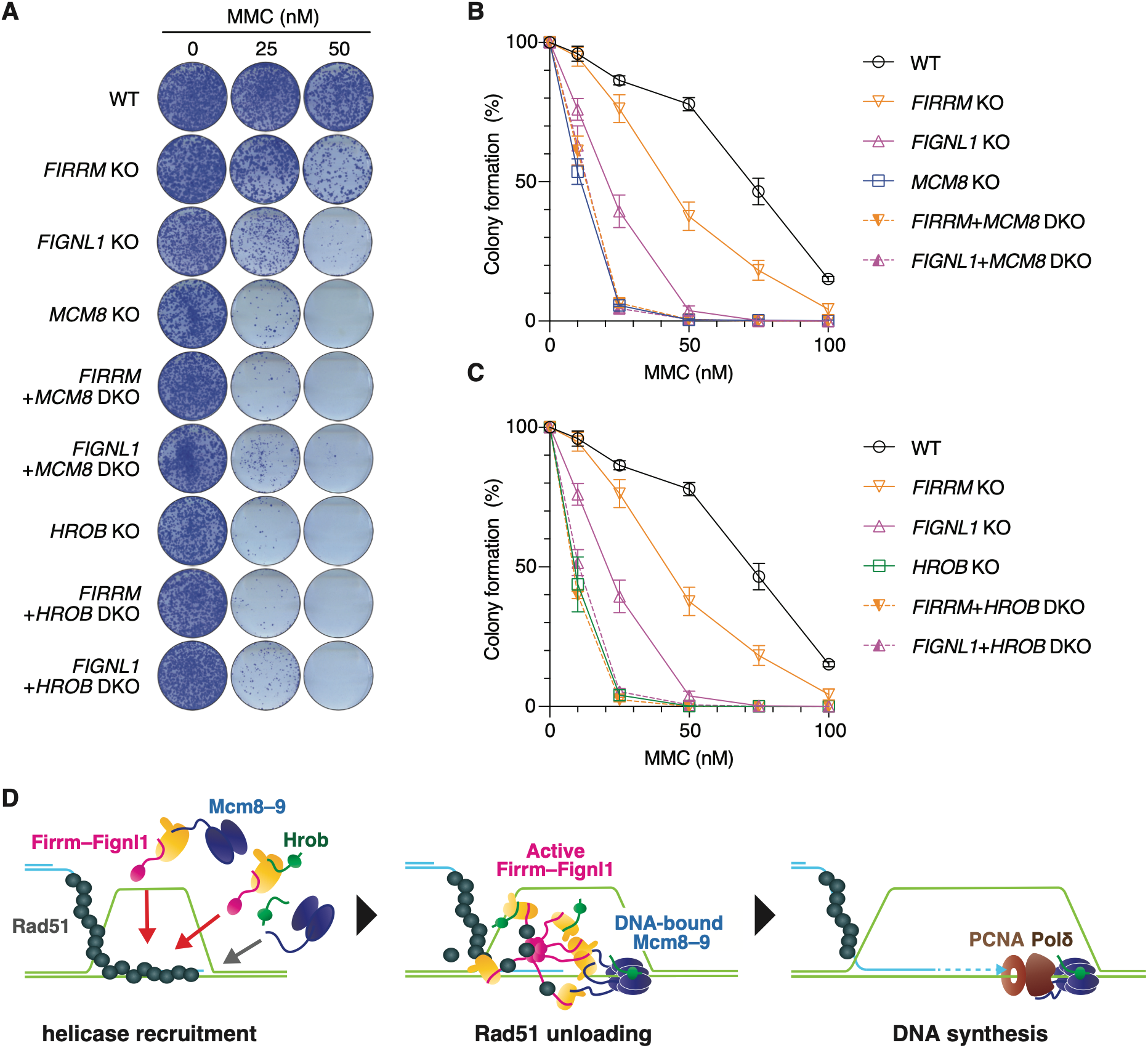
FIRRM–FIGNL1 shows epistasis with MCM8–9 and HROB in the response to mitomycin C. (A) Colony formation assay assessing mitomycin C (MMC) sensitivity in human HCT116 cells of the indicated genotypes. (B, C) Quantification of colony formation by the indicated cells from the three independent experiments. *n* = 3; mean ± SD. (D) Model for the coupling between recombination helicase loading and Rad51 unloading during homologous recombination. See Discussion for details.

Interestingly, *FIGNL1* and *MCM8* double-knockout (*FIGNL1*+*MCM8* DKO) and *FIRRM* and *MCM8* double-knockout (*FIRRM*+*MCM8* DKO) exhibited MMC sensitivity comparable to that of *MCM8* KO cells, indicating that *MCM8* is epistatic to *FIGNL1* and *FIRRM* with respect to MMC resistance (Fig. 7A,B and Supplementary Fig. 9A). Similarly, *HROB* KO showed an epistatic relationship with both *FIGNL1* and *FIRRM* (Fig. 7A,C and Supplementary Fig. 9B). These results indicate that Firrm–Fignl1 and Mcm8–9–Hrob function in the same pathway to confer resistance to DNA interstrand crosslinks in human cells.

## Discussion

Dynamic regulation of Rad51 nucleofilaments is essential for homologous recombination, yet how it is coordinated with downstream recombination events remains poorly understood. In this study, we show that Firrm coordinates the functional interplay between the Fignl1 Rad51 unloader and the Mcm8–9–Hrob recombination helicase, suggesting two principles underlying this coordination. First, the Rad51-unloading machinery promotes recruitment of the recombination helicase to Rad51 filaments, thereby conferring spatial and temporal specificity to helicase recruitment. Second, the recruited recombination helicase reciprocally stimulates the Rad51-unloading machinery, thereby conferring temporal specificity to Rad51 filament disassembly. This reciprocal regulation may represent a molecular ratchet that licenses progression toward recombination-associated DNA synthesis.

Firrm has recently emerged as an important regulator of homologous recombination and Rad51 dynamics (Tsaridou and van Vugt 2024), although its molecular function has remained incompletely understood. Our structural predictions and biochemical analyses show that Firrm directly recognizes short conserved motifs in both Mcm9 and Hrob, providing physical links between Firrm–Fignl1 and the Mcm8–9–Hrob helicase. The Zgrf1 helicase, which facilitates HR by promoting D-loop dissociation, branch migration, and end resection (Brannvoll et al. 2020; Yan et al. 2021), is likewise predicted to interact with Firrm, although through a distinct interface. These findings suggest that Firrm serves as an interaction platform for multiple recombination regulators. Interestingly, previous studies have suggested that Firrm has Fignl1-independent functions that confer resistance to interstrand crosslinking agents (Zhou et al. 2023; Mazouzi et al. 2023; Stok et al. 2023; Pinedo-Carpio et al. 2023). It is therefore tempting to speculate that the interaction platform function of Firrm may, at least in part, operate independently of Fignl1, potentially explaining these observations.

Although our proteomic analysis identified both Rad54B and Fignl1 as Rad51 filament-associated proteins, depletion of Fignl1 demonstrates that it is the primary unloader of Rad51 filaments assembled on dsDNA in *Xenopus* egg extracts. Fignl1 has been proposed to disassemble Rad51 through at least two mechanistically distinct pathways: an ATPase-independent pathway, which employs its Rad51-binding motif (FRBD), and an ATPase-dependent pathway, which involves Fignl1 hexamer formation and its translocation on the Rad51 polypeptides (Matsuzaki et al. 2019; Carver et al. 2025; Kuthethur et al. 2025). Our results identify Mcm8–9 as a regulator of the latter activity. Mcm8–9 stimulated the Fignl1 ATPase in a DNA-dependent manner, and this stimulation required the interaction between Mcm9 and Firrm (Fig. 5). In contrast, Hrob, although able to interact with Firrm, did not stimulate ATP hydrolysis under our assay conditions. These observations argue that Fignl1 activation is not a general consequence of peptide binding to Firrm, but instead requires both Mcm8–9 and DNA. Mcm8–9 forms a heterodimer but assembles into a hexameric ring helicase upon DNA binding (Li et al. 2021; McKinzey et al. 2023; Weng et al. 2023; Acharya et al. 2024; Li et al. 2026). Therefore, one attractive possibility is that DNA-dependent assembly of the Mcm8–9 heterohexamer promotes the recruitment of multiple Firrm–Fignl1 complexes, thereby facilitating assembly of the ATPase-active Fignl1 hexamer responsible for Rad51 unloading (see Fig. 7D). Testing this model will require determination of the higher-order architecture of the Mcm8–9–Firrm–Fignl1 complex, for example by cryo-electron microscopy.

Interestingly, similar phenotypes have been reported for Firrm–Fignl1 and Mcm8–9–Hrob mutants. Independent genetic studies have shown that both complexes contribute to resistance to interstrand crosslinking agents, meiotic recombination, and genome stability (Nishimura et al. 2012; Lutzmann et al. 2012; Lee et al. 2015; Hustedt et al. 2019; Matsuzaki et al. 2019; Olivieri et al. 2020; Wang et al. 2020; Huang et al. 2020; Zhou et al. 2023; Ito et al. 2023; Pinedo-Carpio et al. 2023; Stok et al. 2023; Mazouzi et al. 2023; Zhang et al. 2023; Zainu et al. 2024; Tischler et al. 2024; Cao et al. 2025). Recent studies have further placed both complexes downstream of the Fanconi anemia factors during ICL repair, in which Rad51 plays a critical role (Nishimura et al. 2012; Natsume et al. 2017; Hustedt et al. 2019; Wang et al. 2020; Zhou et al. 2023; Pinedo-Carpio et al. 2023; Mazouzi et al. 2023; Tischler et al. 2024; Beragama Arachchi et al. 2025). Disruption of either complex impairs meiotic crossover formation and leads to persistent accumulation of Rad51- or Dmc1-associated recombination intermediates (Lutzmann et al. 2012; Hustedt et al. 2019; Ito et al. 2023; Zhang et al. 2023; Cao et al. 2025). These striking similarities may be explained, at least in part, by the molecular relationship between Firrm–Fignl1 and Mcm8–9–Hrob uncovered in this study. Indeed, our genetic analyses show that Firrm–Fignl1 and Mcm8–9–Hrob function epistatically in the response to MMC in human cells (Fig. 7A–C). Moreover, a recent genome-wide genetic interaction screen independently placed *FIGNL1* and *MCM8–MCM9–HROB* within the same functional network and predicted a direct interaction between Firrm and Mcm9 using AlphaFold-Multimer (Hayward et al. 2026). This prediction is consistent with an earlier in silico interaction screen (Schmid and Walter 2025), and our analyses provide direct experimental validation of this predicted interaction. Identification of the Firrm-interacting motifs in Mcm9 and Hrob should enable direct dissection of the Firrm-dependent functions of Mcm8–9–Hrob in cellular and genetic contexts.

Collectively, our results support a model in which Firrm–Fignl1 and Mcm8–9–Hrob coordinate the transition from Rad51 unloading to recombination-associated DNA synthesis (Fig. 7D). Firrm–Fignl1 recruits Mcm8–9 and Hrob onto the Rad51–dsDNA filament. Direct interaction between Firrm and Hrob may help stabilize the active Mcm8–9–Hrob helicase at this stage. Mcm8–9 assembles on DNA as a hexameric helicase. This Mcm8–9 hexamer interacts with Firrm and activates the Fignl1 ATPase, possibly by promoting Fignl1 hexamer formation. Fignl1-mediated Rad51 unloading exposes the primer terminus, which is extended by Polδ–PCNA. The interaction between Mcm8–9 and Polδ–PCNA may contribute to the assembly of a recombination-associated DNA synthesis machinery analogous to the CMG–Polε complex at the replication fork (Burgers and Kunkel 2017). Since both Mcm8–9 and Firrm–Fignl1 have been implicated in replication fork protection (Griffin et al. 2022; Zhang et al. 2023; Tischler et al. 2024; Matsuzaki et al. 2024), it will be interesting to examine whether a similar mechanism operates during protection of stalled replication forks. Our study also raises several important mechanistic questions, including where Mcm8–9–Hrob assembles relative to Rad51 filaments, how and when its helicase activity is activated, and whether the Firrm–Mcm9 and Firrm–Hrob interactions have functions beyond helicase recruitment and Fignl1 ATPase activation.

## Methods

### Reagents and resources

Key reagents and resources are provided in Supplementary Table 5. Oligonucleotides are provided in Supplementary Table 6.

### Preparation of *Xenopus* egg extracts

*Xenopus laevis* was purchased from Kato-S-Science and maintained in accordance with the animal experimentation guidelines of Kyushu University. *Xenopus* egg extracts (Walter et al. 1998) were prepared as described previously (Nishiguchi et al. 2025).

### Cloning and plasmids

The *Xenopus laevis rad51, firrm, fignl1, and hrob* coding sequences were amplified from *Xenopus* egg cDNA by PCR using primer pairs 1970/1971, 3006/3007, 3017/3018, and 2363/3489, respectively. The *mcm8* and *mcm9* coding sequences were amplified from pExpress1-xMCM8 and pCMV-SPORT6.ccdb-xMCM9 (Horizon Discovery), respectively, by two-step PCR using primer pairs 1473/1474, followed by 344/345, and 1475/1476, followed by 344/345. All PCR products were cloned into pDONR201 (Thermo Fisher Scientific) by Gateway BP recombination. The resulting Gateway entry clones were designated pDONR-xRAD51, pDONR-xFIRRM, pDONR-xFIGNL1, pDONR-xHROB, pDONR-xMCM8, and pDONR-xMCM9. Epitope tags and protease cleavage sites were introduced into the corresponding Gateway entry clones using the following PCR and cloning strategies: Rad51, 1971/2722, XbaI/SbfI-HF; Fignl1, 3018/3023, followed by 3014/3018, Gateway BP recombination; Hrob, 2363/2411, NdeI/SbfI-HF; and Mcm9, 1490/2600, NdeI/XhoI. For Hrob and Mcm9, pDE1a, a derivative of pDONR201 (Higashi et al. 2012), was used instead of pDONR201. The resulting constructs were designated pDONR-FLAG-TEV-xRAD51, pDONR-FLAG-xFIGNL1, pDE1a-FLAG-xHROB, and pDE1a-FLAG-xMCM9.

Mutations and deletion constructs were generated using the following PCR strategies and restriction enzymes: Fignl1(K428A), 3014/3469 and 3018/3468, overlap-extension PCR with 3014/3018, NcoI-HF/SbfI-HF; Fignl1(1–143), 3023/3491, SacI-HF/SbfI-HF; Mcm8(K453A), 1473/2759 and 1474/2758, overlap-extension PCR with 1473/1474, NcoI-HF/XbaI; Mcm9(K359A), 1491/2761 and 1494/2760, overlap-extension PCR with 1491/1494, NcoI-HF/SacI-HF; Mcm9(1–701), 1491/2932, NcoI-HF/EcoRI-HF; Mcm9(702–1143), 1589/3433, NdeI/EcoRI-HF; Mcm9ΔFIP, 1491/3434, SacI-HF/EcoRI-HF; Mcm9(702–1143ΔFIP), 3433/3434, NdeI/EcoRI-HF; Mcm9(1002–1143), 1589/4039, EcoRI-HF/NdeI; and Mcm9(1002–1143ΔFIP), 3434/4039, EcoRI-HF/NdeI. The resulting constructs were designated pDONR-FLAG-xFIGNL1(K428A), pDONR-FLAG-xFIGNL1(1–143), pDONR-xMCM8(K453A), pDE1a-FLAG-xMCM9(K359A), pDE1a-FLAG-xMCM9(1–701), pDE1a-xMCM9(702–1143), pDE1a-FLAG-xMCM9ΔFIP, pDE1a-xMCM9(702–1143ΔFIP), pDE1a-xMCM9(1002–1143), and pDE1a-xMcm9(1002–1143ΔFIP).

For protein expression in insect cells, coding sequences from the Gateway entry clones were transferred into BaculoDirect C-Term Linear DNA (Thermo Fisher Scientific) by Gateway LR recombination. The following proteins were expressed using this system: Firrm, FLAG-Fignl1, FLAG-Fignl1(K428A), FLAG-Fignl1(1–143), Mcm8, Mcm8(K453A), FLAG-Mcm9, FLAG-Mcm9(K359A), FLAG-Mcm9(1–701), and FLAG-Mcm9ΔFIP. For bacterial protein expression, coding sequences were transferred into Gateway-compatible bacterial expression vectors by Gateway LR recombination. pET21-GHD, a Gateway-compatible derivative of pET21a (Sigma-Aldrich) encoding an N-terminal GST tag, was used to generate pET21-GHD-xMCM9(702–1143), pET21-GHD-xMCM9(702–1143ΔFIP), pET21-GHD-xMCM9(1002–1143), and pET21-GHD-xMCM9(1002–1143ΔFIP). pColdIID, a Gateway-compatible derivative of pColdII (Takara Bio), was used to generate pColdIID-FLAG-TEV-xRAD51, pColdIID-xHROB, and pColdIID-FLAG-xHROB. For bacterial expression as GST fusion proteins, the *hrob* coding sequence was amplified from pDONR-xHROB using primer pair 3617/3620, 3620/3680, 3617/3618, or 3618/3680 and cloned into pGEX-6P-1 (Cytiva) via BamHI-HF and EcoRI-HF digestion to generate pGEX6P1-xHROB, pGEX6P1-xHROBΔFIP, pGEX6P1-xHROB(1–346), and pGEX6P1-xHROB(1–346ΔFIP).

### Protein expression and purification

Rad51, Hrob, and GST fusion proteins were expressed in *Escherichia coli*. The One Shot BL21 Star (DE3) strain (Thermo Fisher Scientific) was used for Rad51, whereas the BL21-CodonPlus(DE3)-RIPL strain (Agilent Technologies) was used for Hrob and GST fusion proteins. Expression from the *cspA* promoter was induced by lowering the temperature from 37°C to 15°C and adding 0.5 mM isopropyl β-D-thiogalactopyranoside (IPTG), and cells were harvested after 24 h. Expression from the T7 promoter was induced by adding 0.5 mM IPTG at 37°C, and cells were harvested after 2 h. For preparation of clarified cell lysates, cells were lysed with 1 mg/mL lysozyme in buffer A1 (50 mM Tris-HCl, pH 8.0, 5% [v/v] glycerol, and 5 mM 2-mercaptoethanol) containing 1× cOmplete EDTA-free protease inhibitor (Roche). The lysate was combined with an equal volume of buffer A2 (50 mM Tris-HCl, pH 8.0, 1 M NaCl, 5% glycerol, 5 mM 2-mercaptoethanol, and 1× cOmplete EDTA-free protease inhibitor), sonicated, and clarified by centrifugation at 67,600 × *g* for 30 min using a Beckman 70Ti rotor. For Hrob and GST fusion proteins, buffers B1 (30 mM Tris-HCl, pH 8.0, 1 mg/mL lysozyme) and B2 (100 mM Na-phosphate, 1 M NaCl, 2% [w/v] Triton X-100, 10 mM 2-mercaptoethanol, 2 mM phenylmethylsulfonyl fluoride [PMSF]) were used in place of buffers A1 and A2, respectively. For GST fusion proteins, clarification was performed at 20,000 × *g* using a benchtop microcentrifuge instead of a Beckman 70Ti rotor.

For Rad51 purification, the clarified cell lysate was passed through a DEAE Sepharose Fast Flow column (Cytiva), and the flow-through was loaded onto a FLAG-M2 agarose column (Sigma Aldrich). The column was washed with buffer A3 (50 mM Tris-HCl, pH 7.4, 1 mM DTT, 100 mM NaCl, and 5% glycerol), and bound proteins were eluted by on-column cleavage with TEV protease in buffer A3 overnight at 4°C. Following TEV elution, residual bound proteins were recovered by washing the column with buffer A4 (50 mM Tris-HCl, pH 8.0, 5% glycerol, 1% Triton X-100, 500 mM NaCl, 5 mM 2-mercaptoethanol). The eluates were pooled and diluted with buffer A1 to a final NaCl concentration of 100 mM, and loaded onto a MonoQ 5/50 GL column (Cytiva, #17516601). Bound proteins were eluted using a 0.1–1 M linear NaCl gradient in buffer A5 (20 mM Tris-HCl, pH7.4, 10% glycerol, and 5 mM 2-mercaptoethanol). Peak fractions were pooled and dialyzed against buffer A6 (20 mM Tris-HCl, pH 7.4, 150 mM NaCl, 10% glycerol, and 5 mM 2-mercaptoethanol). Aliquots were snap-frozen in liquid nitrogen and stored at −80°C.

For Hrob purification, the clarified cell lysate was incubated with TALON Metal Affinity Resin (Clontech) at 4°C for 90 min with gentle rotation. The resin was transferred to an open-top column, and bound proteins were eluted with buffer B3 (50 mM Na-phosphate, 500 mM NaCl, 1% Triton X-100, 5 mM 2-mercaptoethanol, 1 mM PMSF) containing 0.2 M imidazole. Peak fractions were pooled and diluted fourfold with buffer C (25 mM Tris-HCl, pH 7.4, 5 mM 2-mercaptoethanol, 2.5 mM MgCl_2_, 10% glycerol), loaded onto a HiTrap Q HP column (Cytiva), and bound proteins were eluted using a 0.1–1 M linear NaCl gradient in buffer C. Peak fractions were pooled and subjected to size-exclusion chromatography on a Superdex 200 Increase 10/300 GL column (Cytiva) equilibrated with buffer C containing 150 mM NaCl. Protein-containing fractions were pooled, aliquoted, snap-frozen in liquid nitrogen, and stored at −80°C.

Mcm8–9, Firrm–Fignl1, and their mutant complexes were expressed in Sf9 insect cells by co-infection with baculoviruses encoding the corresponding proteins. Cells were cultured at 28°C in Sf-900 II SFM medium (Thermo Fisher Scientific) supplemented with 2% (v/v) fetal bovine serum (FBS), harvested, washed with phosphate-buffered saline (PBS), snap-frozen in liquid nitrogen, and stored at −80°C. For preparation of clarified cell lysates, cells were suspended in buffer C containing 500 mM NaCl, 1 mM PMSF, and 1× cOmplete EDTA-free protease inhibitor. The cell lysate was clarified by centrifugation at 81,800 × *g* for 30 min using a Beckman 70Ti rotor (Beckman Coulter) and passed through a DEAE Sepharose Fast Flow column (Cytiva) to remove nucleic acids.

For purification of Mcm8–9 and its mutant complexes, the clarified cell lysate was applied to a FLAG-M2 agarose column, and bound proteins were eluted with buffer C containing 500 mM NaCl, and 50 μg/mL FLAG-peptide (Sigma Aldrich). Peak fractions were pooled, diluted 3.3-fold with buffer C, and loaded onto a MonoQ 5/50 GL column (Cytiva). Bound proteins were eluted using a 0.15–1 M linear NaCl gradient in buffer C. Peak fractions were pooled and dialyzed against buffer C containing 150 mM NaCl. Aliquots were snap-frozen in liquid nitrogen and stored at −80°C.

For purification of Firrm–Fignl1 and its mutant complexes, FLAG affinity purification was performed as described for Mcm8–9. The eluates were pooled and concentrated using Amicon Ultra centrifugal filter (0.5 mL or 4.0 mL) (Merck). Concentrated samples were layered onto a 4.5 mL 15%–35% (v/v) glycerol gradient over a 500-μL 50 % (v/v) glycerol cushion prepared with buffer D (25 mM Tris-HCl pH 7.4, 150 mM NaCl, 5 mM 2-mercaptoethanol, and 2.5 mM MgCl_2_) and centrifuged at 81,800 × *g* for 14 h using a Beckman SW55Ti rotor (Beckman Coulter). Gradient fractions were collected, aliquoted, snap-frozen in liquid nitrogen, and stored at −80°C.

### Immunological methods

Antibodies used in this study are listed in Supplementary Table 5. Rabbit antisera against *Xenopus* Mcm8, Mcm9, Hrob, Firrm, and Fignl1 were raised against each N-terminally His6-tagged recombinant protein expressed in *E. coli*. For immunoblotting, Mcm8, Mcm9, Hrob, Firrm, and Fignl1 antisera were used at 1:5,000, and Rad51, Pcna, Msh2, and Msh6 antisera were used at 1:10,000. Alexa Fluor 647/680-conjugated Goat anti-rabbit IgG (H+L) antibodies or Alexa Fluor 647/680-conjugated Mouse anti-Rabbit IgG (light chain specific) (Jackson ImmunoResearch) were used at 1:10,000 as the secondary antibodies.

For immunodepletion, 3 volumes of an antiserum were coupled to 1 volume of recombinant Protein A Sepharose Fast Flow (PAS; Cytiva) at 4°C overnight. 0.2 volumes of the antibody-coupled PAS beads were incubated with 1 volume of *Xenopus* egg extract at 4°C for 1 h, and the procedure was repeated twice. Preimmune serum was used for mock depletion. Typically, 10–90 μL of extract was used per assay.

For immunoprecipitation, 0.5 volumes of antiserum were coupled to 1 volume of recombinant PAS. Five volumes of *Xenopus* egg extract diluted 10-fold with ELB (10 mM Hepes-KOH, 2.5 mM MgCl_2_, 50 mM KCl, pH 7.7) were added to 1 volume of IgG-coupled PAS beads, and the mixture was incubated at 4°C for 2 h with gentle rotation. Typically, 50–100 μL of diluted extract was used per sample. The supernatant and beads were separated by centrifugation at 2,200 × *g* for 1 min using a TMS-21 swinging-bucket rotor (Tomy Seiko). The beads were washed three times with 200 μL of ELB and resuspended in Laemmli SDS sample buffer (62.5 mM Tris-HCl, 10% glycerol, 3% SDS, 0.005% bromophenol blue, 5% 2-mercaptoethanol, pH 6.8).

### Recombinant protein pull-down assay

Anti-FLAG-M2 agarose beads (5 μL per sample) were washed with ELB, mixed with 5 μg of FLAG-tagged recombinant proteins, incubated for 2 h with gentle rotation, and washed three times with 200 μL of ELB. A 4-μL aliquot of the beads was then incubated with 36 μL of *Xenopus* egg extract diluted 5-fold with ELB for 1 h at 4°C with gentle rotation. The beads were washed three times with 200 μL of ELB, and bound proteins were eluted with Laemmli SDS sample buffer.

### Glycerol gradient sedimentation

A 40-μL aliquot of *Xenopus* egg extract was layered onto a 4.5-mL 15%–35% (v/v) glycerol gradient formed over a 500-μL 50% glycerol cushion, using ELB and ELB containing 50% glycerol as the light and heavy solutions, respectively. Gradients were centrifuged at 81,800 × *g* for 14 h at 4°C in a Beckman SW55Ti rotor. Following centrifugation, 24 fractions (approximately 200 μL each) were manually collected from the bottom of the gradient, mixed with Laemmli SDS sample buffer, and analyzed by SDS-PAGE and immunoblotting. The gradient was calibrated in parallel using molecular mass standards (thyroglobulin, 669 kDa; apoferritin, 443 kDa; catalase, 232 kDa; bovine serum albumin, 67 kDa; and carbonic anhydrase, 25 kDa).

### GST pull-down assay

Glutathione Sepharose 4B (5 μL per sample) was washed with buffer B3, mixed with 500 μL of lysates from *E. coli* cells expressing GST-tagged recombinant proteins, incubated at 4°C for 2 h with gentle rotation, and washed three times with 100 μL of buffer C containing 150 mM NaCl and 1% Triton X-100. A 4-μL aliquot of the beads was then incubated with 36 μL of 100 nM Firrm–Fignl1(1–143) in buffer C containing 150 mM NaCl and 1% Triton X-100 for 1h at 4°C with gentle rotation. The beads were washed three times with 100 μL of buffer C containing 150 mM NaCl and 1% Triton X-100, and bound proteins were eluted with Laemmli SDS sample buffer.

### DNA substrate preparation

Poly-dA/dT DNA for the ATPase assay was prepared by mixing 10 μM each of 70-mer poly-dA and poly-dT oligonucleotides in TEN buffer (10 mM Tris–HCl, pH 7.4, 1 mM EDTA, 100 mM NaCl), heating the mixture at 95°C for 5 min, and allowing it to anneal by incubation at 25°C for 10 min. Biotinylated DNA substrates used for the Rad51 filament pull-down assay were prepared essentially as described previously (Shigenobu-Ueno et al. 2025). Briefly, 40 μg of pMMR1 plasmid (Shigenobu-Ueno et al. 2025) was nicked with 100 units of Nb.BtsI (NEB) in 400 μL of 1× rCutSmart buffer (NEB) for 2 h at 37°C. The DNA fragments flanked by the nicks were then replaced with the 5′-phosphorylated oligonucleotide 2983 containing two biotinylated thymines, ligated, and purified. The resulting biotinylated DNA was immobilized on Sepharose beads at a ratio of 100 ng DNA per 1 μL of beads using the procedure described previously (Higashi et al. 2012).

### ATPase assay

ATPase reactions (20 μL) containing 50 nM of each recombinant protein in buffer D (25 mM Tris-HCl, pH 7.4, 25 mM NaAc, 1 mM MgCl_2_, 1 mM DTT, 0.1 mg/mL BSA, 0.6 mM ATP) were incubated at 22°C for 2 h. Where appropriate, poly-dA/dT DNA was added at a final concentration of 1 μM. Reactions were then mixed with 10 μL of H_2_O and 70 μL of CytoPhos reagent (Cytoskeleton), and inorganic phosphate release was quantified by measuring absorbance at 650 nm using a Multiskan FC microplate reader (Thermo Fisher Scientific).

### Rad51 filament pull-down assay

To assemble Rad51 filaments, 1 volume of DNA beads was washed with and resuspended in 2.5 volumes of buffer E (25 mM Tris-HCl, pH 7.4, 5 mM MgCl_2_, 1 mM DTT, 100 ng/μL BSA, 0.1% Triton X-100, 1 mM ATP, 2 mM CaCl_2_) containing Rad51 at the desired concentrations, and incubated at 22°C for 30 min. The beads were then washed with 500 μL ELB once and incubated at 22°C in *Xenopus* egg extracts supplemented with 2 mM ATP, 20 mM phosphocreatine (PC), and 5 μg/mL creatine phosphokinase (CPK), which had been prewarmed at 22°C for 10 min. Typically, 1.5–2.2 μL of Sepharose beads were incubated in 6–8.8 μL of extract, corresponding to a final concentration of 20 ng/μL DNA. At the appropriate time points, the mixture was quickly diluted with 200 μL of ELB containing 0.1% Triton X-100, layered onto a 200-μL ELB cushion containing 0.5 M sucrose, and centrifuged at 12,700 × *g* for 1 min at 4°C in a TMS-21 swinging bucket rotor. DNA beads were then divided into two aliquots for analysis of DNA-binding factors and bead-bound DNA. To analyze DNA-binding proteins, one aliquot was mixed with Laemmli SDS sample buffer, and the bead-bound proteins were analyzed by SDS-PAGE followed by immunoblotting or mass spectrometry. For DNA quantification, the other aliquot was mixed with 100 μL of stop buffer (1% sodium dodecyl sulfate [SDS], 20 mM EDTA, 3 mM biotin) and treated with 50 μg/mL Proteinase K (Nacalai Tesque) at 37°C overnight. DNA was purified by phenol/chloroform extraction and ethanol precipitation, resuspended in TE buffer (10 mM Tris-HCl pH 7.4, 1 mM EDTA) containing 10 μg/mL RNase A, digested with EcoRI-HF, and analyzed by 0.8% agarose gel electrophoresis followed by SYBR Gold staining (Thermo Fisher Scientific).

### LC-MS/MS analysis

Mass spectrometry samples were prepared as described previously (Miyata et al. 2026). Peptides were analyzed using either an Advance UHPLC system (Michrom Bioresources) coupled to a Q Exactive mass spectrometer (Thermo Fisher Scientific) or an Evosep One system (EvoSep Biosystems) coupled to an Orbitrap Excedion Pro mass spectrometer (Thermo Fisher Scientific). For the Evosep system, peptides were separated on an Aurora Elite column (15 cm × 75 μm ID, IonOpticks) using the Whisper Zoom 20 SPD method. Database searching and data validation were performed as described previously (Miyata et al. 2026).

### Mass spectrometry data analysis

Peptides were identified using Scaffold version 5 (Proteome Software), and spectral counts (Total Spectra) were used for quantitative analysis. Proteins detected in at least 3 out of 6 samples (2 conditions with 3 biological replicates each) were subjected to further analysis. The data were log_2_-transformed prior to statistical analysis, and missing values were imputed by random draws from a Gaussian distribution N(μ − 1.5σ, (0.3σ) ²), where μ and σ are the mean and standard deviation of the observed log_2_-transformed values, under the assumption that missing values predominantly arise from low-abundance proteins (Tyanova et al. 2016). Log_2_ fold changes were calculated between conditions, and statistical significance was assessed using paired *t*-tests.

### AlphaFold3 structural modeling

Complex structures were predicted using AlphaFold3 (Abramson et al. 2024) through the AlphaFold Server. Models were generated with 10 recycles (ranking score: Firrm–Fignl1N–Mcm9, 0.60; Firrm–Fignl1N–Hrob, 0.76; Firrm–Fignl1–Zgrf1, 0.48). Predicted structures were visualized in ChimeraX (Pettersen et al. 2021), and electrostatic surface potentials were calculated using the Coulombic surface coloring tool implemented in ChimeraX. Predicted aligned error (PAE) matrices generated by AlphaFold3 were visualized as heat maps using Matplotlib.

### Cell culture and gene knockout

HCT116 cells were cultured in McCoy’s 5A, supplemented with 10% FBS, 2 mM L-glutamine, 100 U/mL penicillin and 100 μg/mL streptomycin at 37 °C in a 5% CO_2_ incubator. The plasmids used for gene knockout are listed in Supplementary Table 5. To generate *MCM8* KO cell lines, HCT116 cells were transfected with plasmids expressing Cas9 and a gRNA targeting exon 7 of the *MCM8* gene (Supplementary Table 5, 7). Following transfection, individual clones were isolated and analyzed by sequencing to confirm frameshift mutations, and the loss of Mcm8 protein was confirmed by immunoblotting. To knock out *FIRRM*, *FIGNL1* and *HROB*, we designed CRISPR plasmids coexpressing a gRNA and Cas9, together with donor plasmids carrying a puromycin- or blasticidin-resistant marker (Supplementary Table 5, 7). HCT116 cells were transfected with the CRISPR-Cas9 and donor plasmids and selected according to a previously published protocol (Saito and Kanemaki 2021). Individual clones were isolated following selection with 1 μg/mL puromycin or 10 μg/mL blasticidin S. Clones harboring bi-allelic insertions at the target locus were identified by genomic PCR, and the loss of the target protein was confirmed by immunoblotting.

### Colony formation assay

Cells were seeded in 6-well plates at a density of 3,000 cells pre well. One day after seeding, cells were treated with mitomycin C (MMC, Kyowa Kirin) at concentrations of 0, 10, 25, 50, 75, or 100 nM. After 24 h of treatment, the medium was replaced with fresh medium without MMC, and the cells were cultured for an additional week. Cells were then fixed and stained with crystal violet solution (6.0% glutaraldehyde and 0.5% crystal violet). Images of the stained cell colonies were analyzed using Fiji, and the total colony area value per well was quantified. For each experiment, the value for the untreated cells was set to 100%, and all other values were expressed relative to this value. All experiments were performed three times independently.

### Statistical analyses

Statistical analyses were performed using Prism 10.1 (GraphPad Software). Details of the individual statistical tests are indicated in the figure legends and results.

### Generative AI

Generative AI was used during manuscript preparation to assist with language editing. It was not used to generate, analyze, or interpret experimental data. The authors reviewed and revised all AI-assisted text and take full responsibility for the final manuscript.

## Data availability

All data supporting the findings of this study are available within the paper and its Supplementary Information. Unprocessed raw data of mass spectrometry are available at https://doi.org/10.48708/7441980. Materials generated in this study are available from the corresponding authors upon reasonable request.

## Acknowledgments

We thank Satomi Oda for assistance in the preparation of Mcm8 antisera, Shota Fukuda for assistance in the preparation of Rad51 antisera, Toyoaki Natsume (Tokyo Metropolitan Institute of Medical Science) for providing *MCM8* KO cells, and the members of the Takahashi and Kanemaki laboratories for helpful discussions.

## Author contributions

D.N. performed most of the *Xenopus* experiments. Y.T. and M.T.K. performed the human cell experiments. N.T. and K.I. performed the mass spectrometry analyses. K.T. optimized the Rad51 loading conditions. Y.N. and Y.K. contributed to experimental design, assisted with experiments, and participated in data interpretation. D.N., M.T.K., and T.S.T. conceived and designed the study and analyzed the data. D.N. drafted the manuscript. M.T.K. and T.S.T. revised and edited the manuscript. All authors reviewed and approved the final manuscript.

