## Supplemental Figures and Legends for "Firrm functions as a multiprotein interaction platform that coordinates Mcm8–9–Hrob recruitment and Fignl1-mediated Rad51 unloading"

#### Supplementary Figure 1

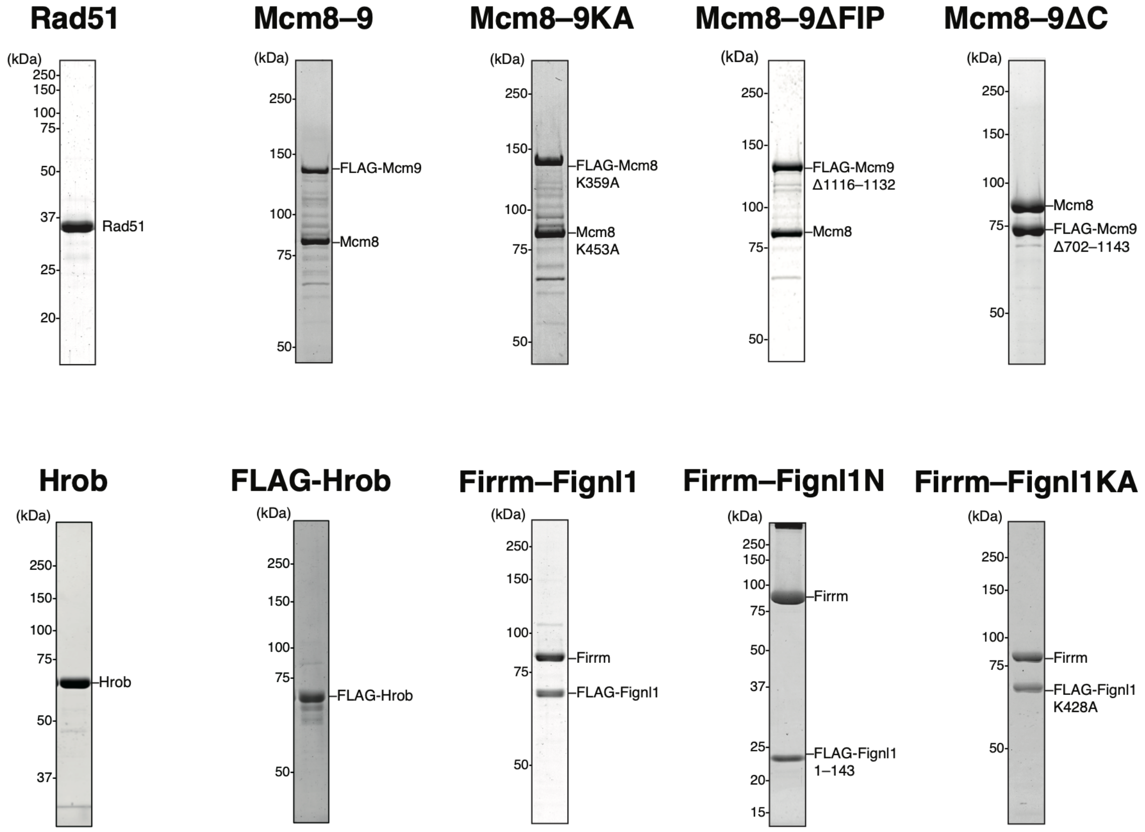


**Supplementary Figure 1. Recombinant proteins used in this study.**

Recombinant proteins were separated by SDS-PAGE and stained with Coomassie Brilliant Blue R-250. Rad51, wild-type Rad51; Mcm8–9, wild-type Mcm8/FLAG-Mcm9; Mcm8–9KA, Mcm8(K453A)/FLAG-Mcm9(K359A); Mcm8–9ΔFIP, wild-type Mcm8/FLAG-Mcm9(Δ1116–1132); Mcm8–9ΔC, wild-type Mcm8/FLAG-Mcm9(Δ702–1143); Hrob, wild–type Hrob; FLAG-Hrob, FLAG-tagged Hrob; Firrm–Fignl1, wild-type Firrm/FLAG-Fignl1; Firrm–Fignl1N, wild-type Firrm/FLAG-Fignl1(1–143); Firrm–Fignl1KA, wild-type Firrm/FLAG-Fignl1(K428A).

### Supplementary Figure 2

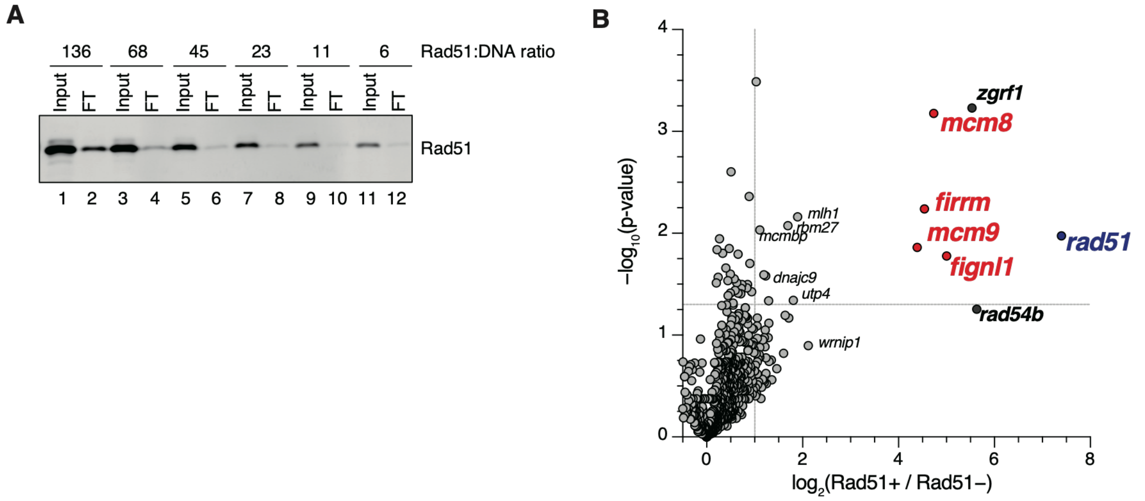


**Supplementary Figure 2. Proteomic analysis of proteins associated with Rad51 filaments assembled on dsDNA.**

(A) Rad51 filaments were assembled on DNA beads at the indicated Rad51:DNA ratios. Input and flow-through (FT) fractions were analyzed by immunoblotting.

(B) Volcano plot of $\log_2$ fold change versus $-\log_{10}(P\text{ value})$ for proteins identified in pull-downs performed with and without Rad51 filaments. Dashed lines indicate $P = 0.05$ and a $\log_2$ fold change of 1.

#### Supplementary Figure 3

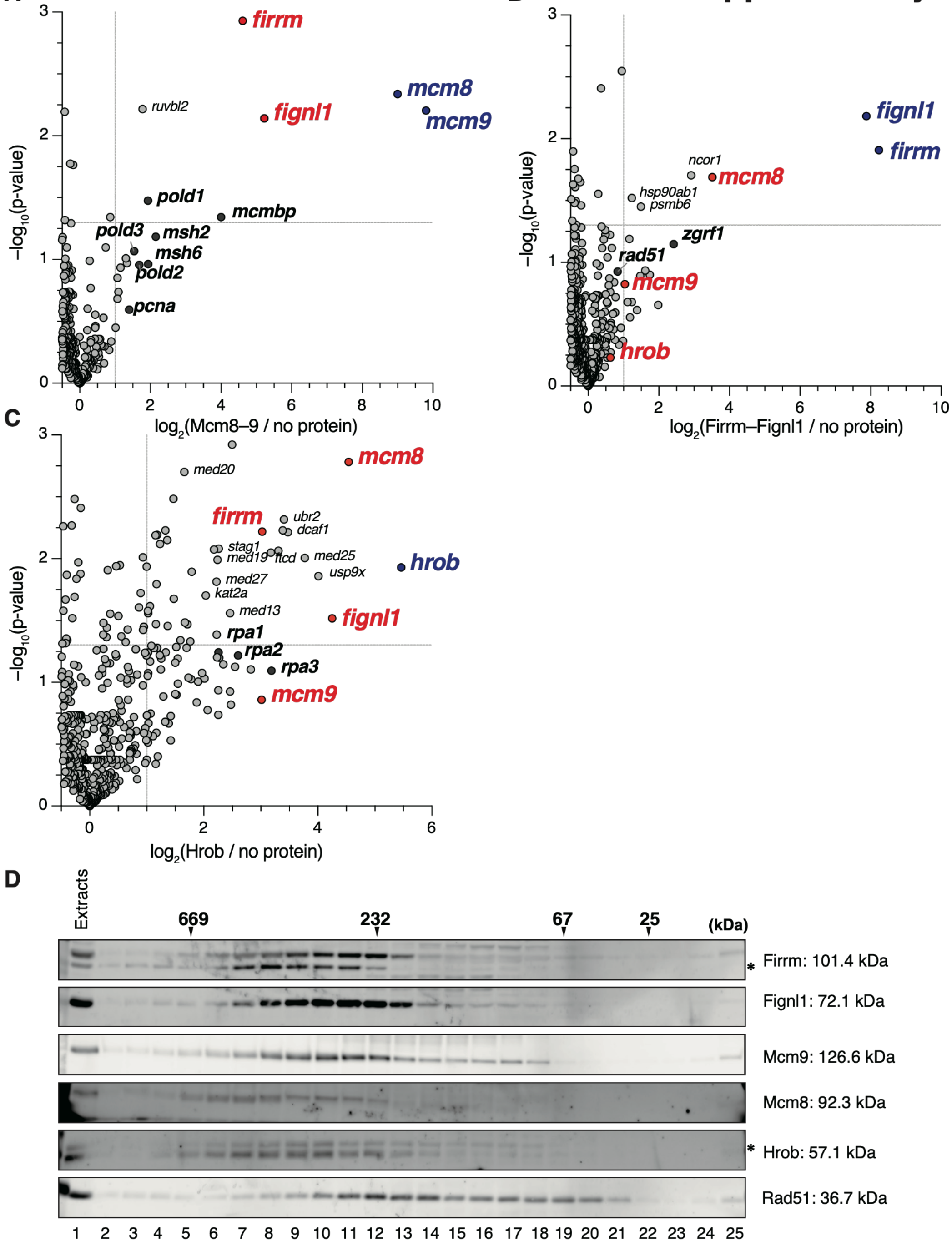


**Supplementary Figure 3. Interaction analyses of Firrm–Fignl1, Mcm8–9, and Hrob in *Xenopus* egg extracts.**

(A–C) Volcano plots of $\log_2$ fold change versus $-\log_{10}$($P$ value) for proteins identified in pull-downs with Mcm8–9 (A), Firrm–Fignl1 (B), or Hrob (C), each compared with the no-protein control. Dashed lines indicate $P = 0.05$ and a $\log_2$ fold change of 1.

(D) Glycerol gradient sedimentation of *Xenopus* egg extract followed by immunoblot analysis. Asterisks indicate cross-reacting bands. Molecular mass standards: thyroglobulin (669 kDa), catalase (232 kDa), albumin (67 kDa), and chymotrypsinogen A (25 kDa).

#### Supplementary Figure 4

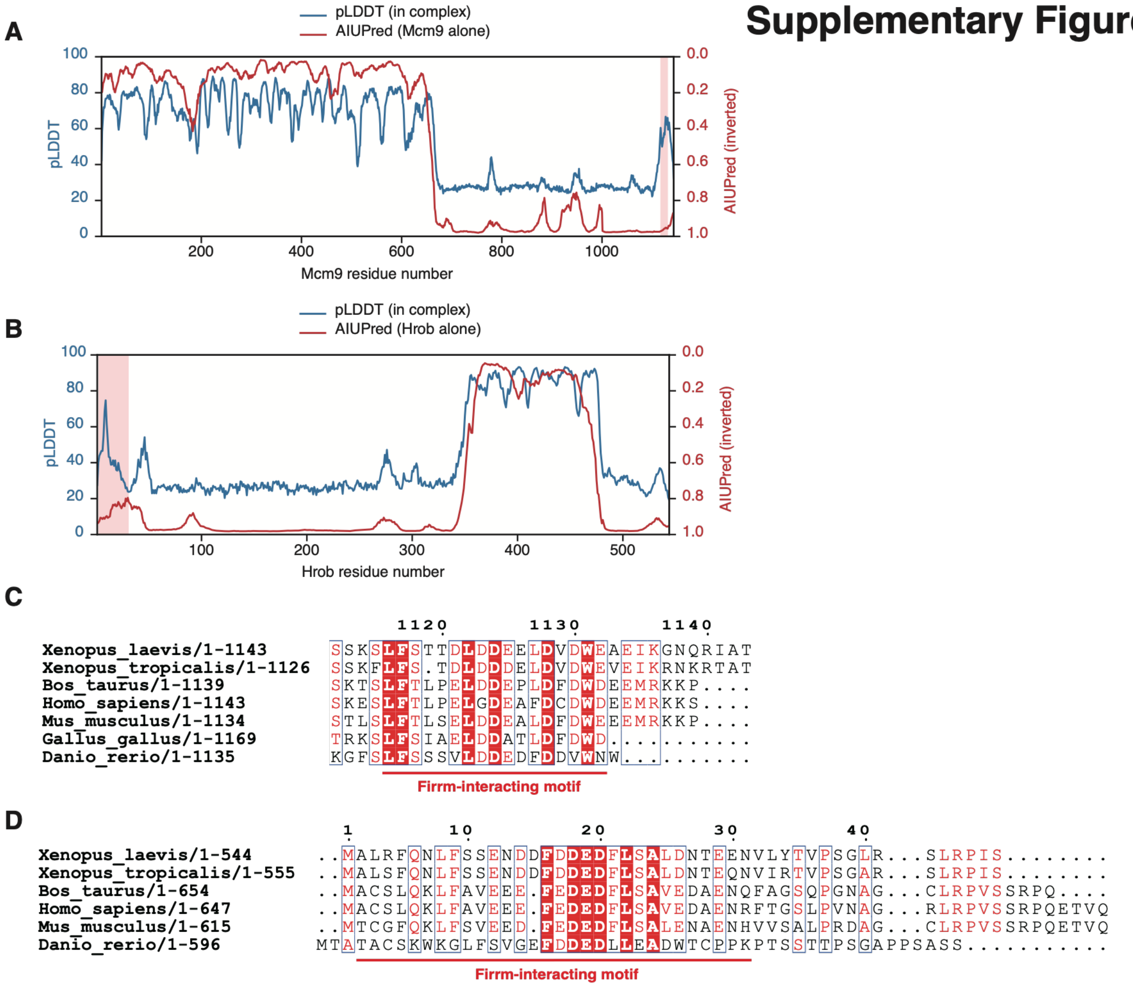


**Supplementary Figure 4. Firrm-interacting motifs in Mcm9 and Hrob are conserved across species and reside within intrinsically disordered regions.**

(A, B) Intrinsic disorder propensity of Mcm9 (A) and Hrob (B) was predicted using AIUPred (Erdős et al. 2021; https://aiupred.elte.hu) with default smoothing settings. The predicted disorder profiles are compared with the AlphaFold3 per-residue pLDDT values.

(C, D) Orthologous Mcm9 (C) and Hrob (D) protein sequences from *Homo sapiens*, *Mus musculus*, *Gallus gallus*, *Xenopus tropicalis*, *Bos taurus*, and *Danio rerio* were aligned using ClustalW (Thompson et al. 1994) and visualized using ESPript 3 (Robert and Gouet 2014; https://espript.ibcp.fr)

### Supplementary Figure 5

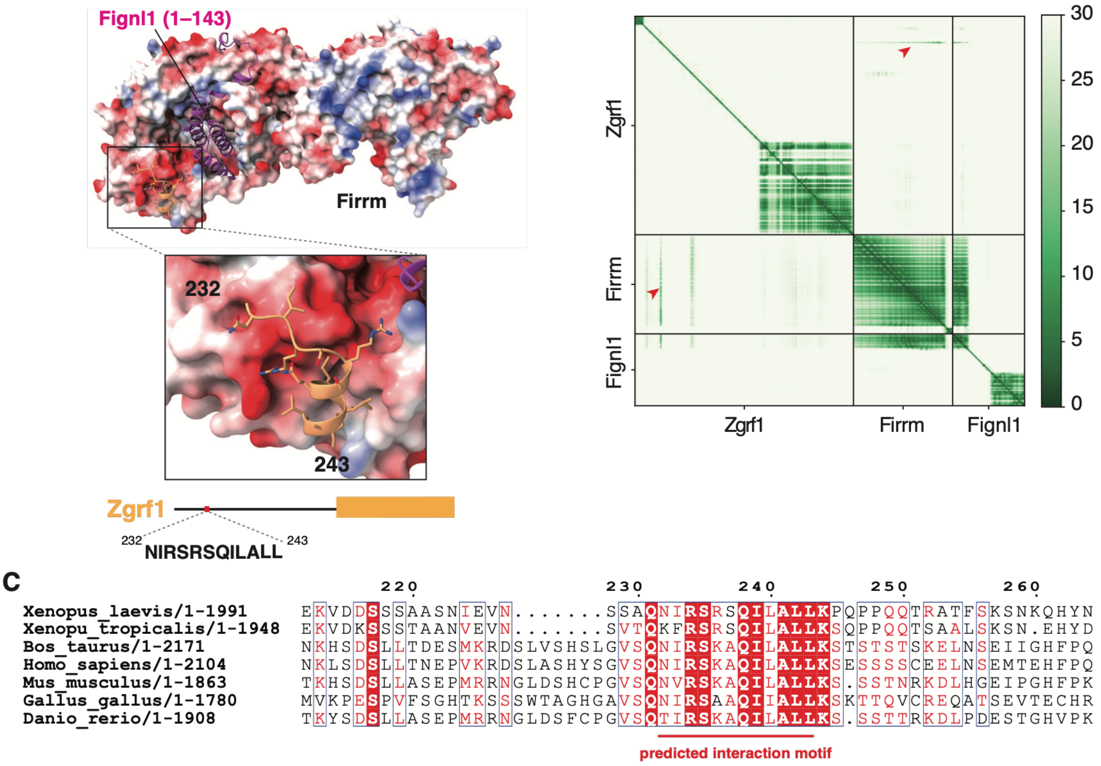


**Supplementary Figure 5. A conserved Zgrf1 motif is predicted to interact with Firrm.**

(A) AlphaFold3-predicted structure of the Firrm–Fignl1 complex bound to a Zgrf1-derived peptide. The surface of Firrm is colored according to its electrostatic surface potential, with basic and acidic regions shown in blue and red, respectively. The Zgrf1-derived peptide is shown in yellow. The residue range and amino acid sequence of the peptide are shown below.

(B) Predicted aligned error (PAE) heat map for the AlphaFold3 model shown in (A). Red arrowheads indicate the predicted Firrm–Zgrf1 interface.

(C) Sequence alignment of Zgrf1 across species. The predicted Firrm-interacting motif is underlined.

### Supplementary Figure 6

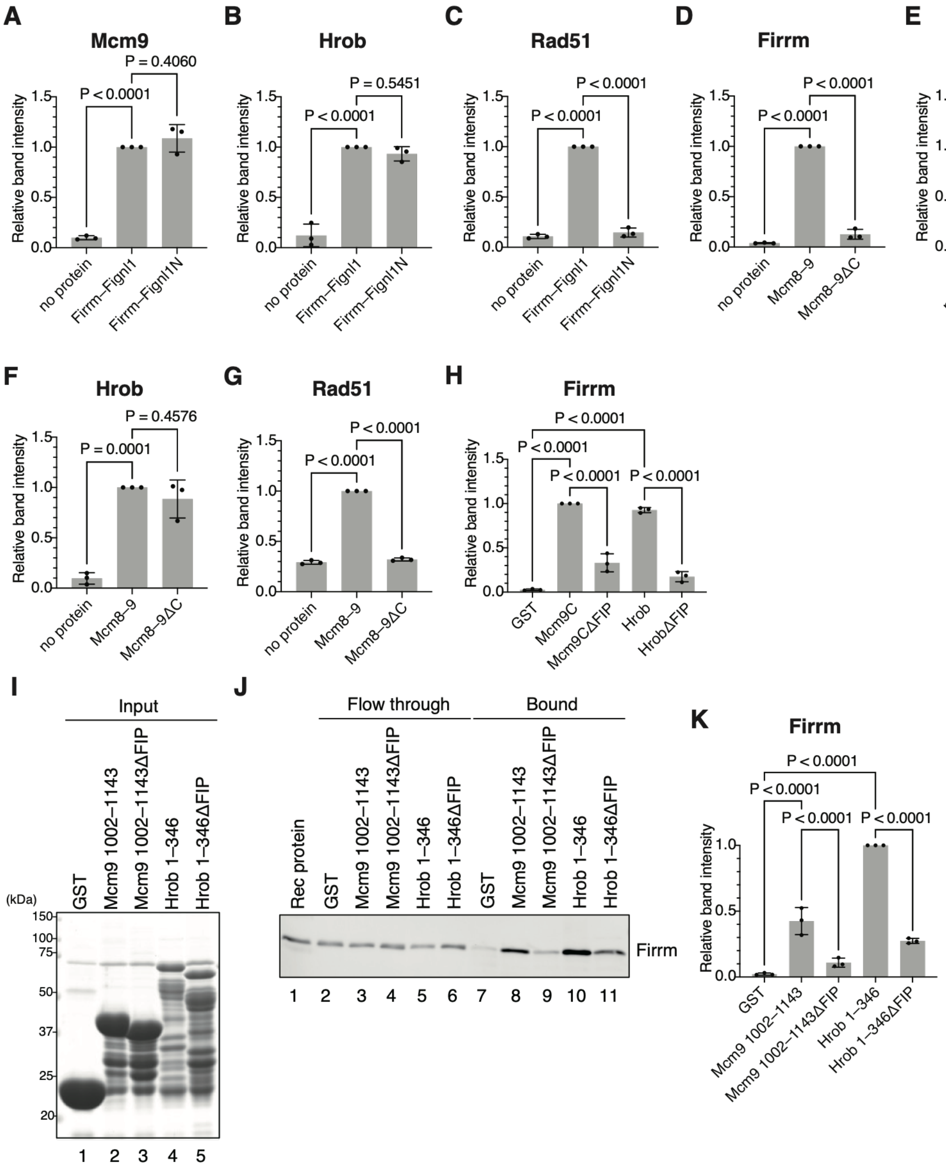


**Supplementary Figure 6. Firrm functions as a multiprotein interaction platform.**

(A–C) Relative band intensities of Mcm9 (A), Hrob (B), and Rad51 (C) were calculated from replicate experiments in Fig. 3D, with the signal in lane 6 set to 1. $n = 3$; mean ± SD; one-way ANOVA with Šídák's multiple comparisons test.

(D–G) Relative band intensities of Firrm (D), Msh6 (E), Hrob (F), and Rad51 (G) were calculated from replicate experiments in Fig. 3F, with the signal in lane 6 set to 1. $n = 3$; mean ± SD; one-way ANOVA with Šídák's multiple comparisons test.

(H) The relative band intensities of Firrm were calculated from replicate experiments in Fig. 3H, with the signal in lane 8 set to 1. $n = 3$; mean ± SD; one-way ANOVA with Šídák's multiple comparisons test.

(I, J) Recombinant pull-down assay using GST, GST-Mcm9-1002–1143, GST-Mcm9-1002–1143ΔFIP [Mcm9(1002–1143, Δ1116–1132)], GST-Hrob-1–346, and GST-Hrob 1–346ΔFIP [Hrob(1–346, Δ2–31)]. GST fusion proteins were immobilized on Glutathione Sepharose beads, incubated with recombinant Firrm–Fignl1N, and recovered. Coomassie Brilliant Blue R-250 staining of input samples (I) and immunoblot analysis of flow-through and bound fractions probed with Firrm antibodies (J) are shown. A representative experiment from three technical replicates is shown.

(K) The relative band intensities of Firrm were calculated from replicate experiments in (J), with the signal in lane 10 set to 1. $n = 3$; mean ± SD; one-way ANOVA with Šídák's multiple comparisons test.

#### Supplementary Figure 7

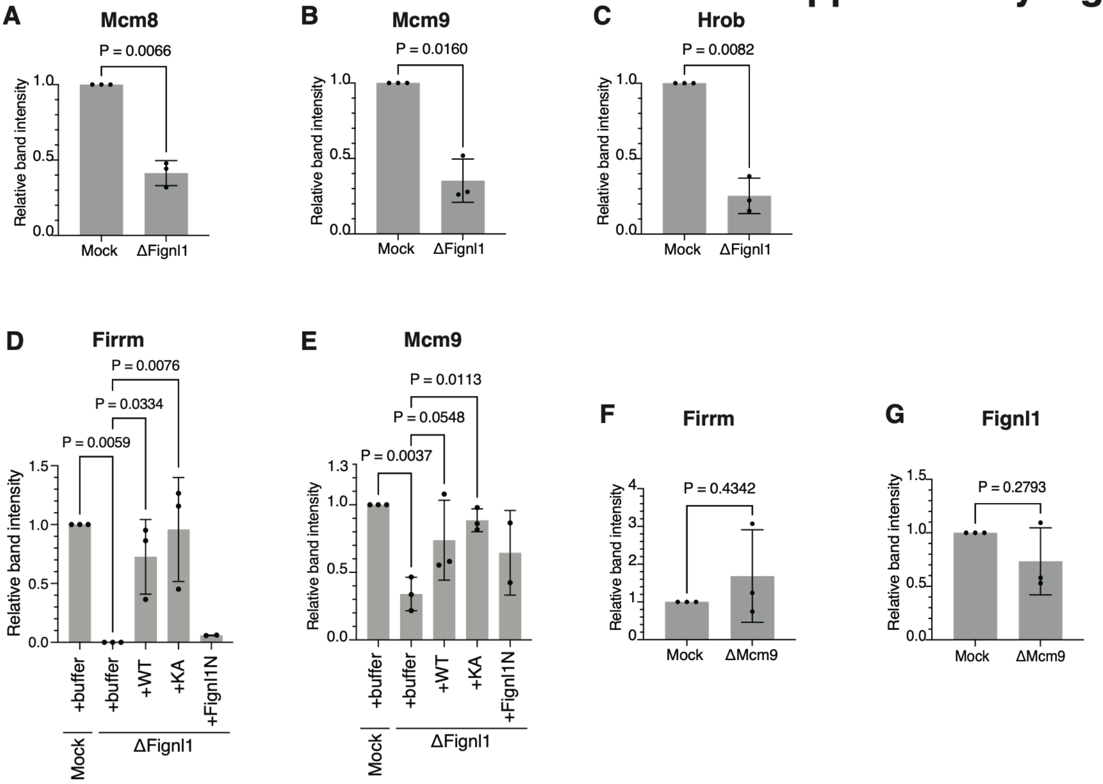


**Supplementary Figure 7. Firrm–Fignl1 promotes Mcm8–9–Hrob recruitment to Rad51 filaments through the Firrm–Mcm9 interaction.**

(A–C) Relative band intensities of Mcm8 (A), Mcm9 (B), and Hrob (C) were calculated from the 30-min time point of replicate experiments in Fig. 4B, with the signal in lane 4 set to 1. $n = 3$; mean ± SD; paired *t*-test

(D, E) Relative band intensities of Firrm (D) and Mcm9 (E) were calculated from replicate experiments in Fig. 4D, with the signal in lane 2 set to 1. $n = 3$; mean ± SD; one-way ANOVA with Šídák's multiple comparisons test.

(F, G) Relative band intensities of Firrm (F), and Fignl1 (G) were calculated from the 30-min time point of replicate experiments in Fig. 4H, with the signal in lane 4 set to 1. $n = 3$; mean ± SD; paired *t*-test

### Supplementary Figure 8

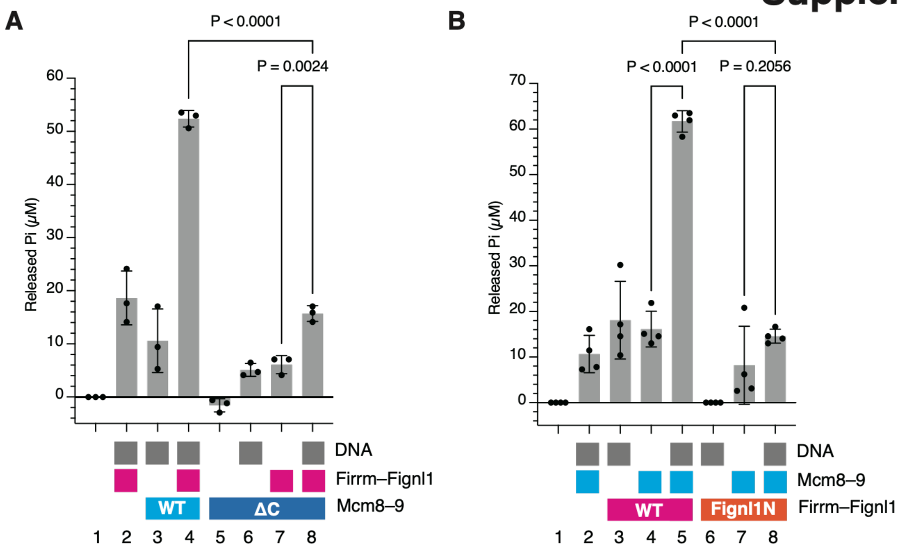


**Supplementary Figure 8. Mcm8–9-dependent stimulation of Fignl1 ATPase activity requires the C-terminal IDR of Mcm9 and the C-terminal region of Fignl1.**

(A, B) ATPase activity was quantified by measuring released inorganic phosphate. Recombinant proteins were used at 50 nM each: Firrm–Fignl1, Mcm8–9, and Mcm8–9ΔC [Mcm9(Δ702–1143)] in (A); Mcm8–9, Firrm–Fignl1, and Firrm–Fignl1N [Fignl1(1–143)] in (B). Poly-dA/dT DNA was added at 1 µM. Squares indicate the proteins and DNA present in each reaction. *n* = 3 (A) or *n* = 4 (B); mean ± SD; one-way ANOVA with Šídák's multiple comparisons test.

#### Supplementary Figure 9

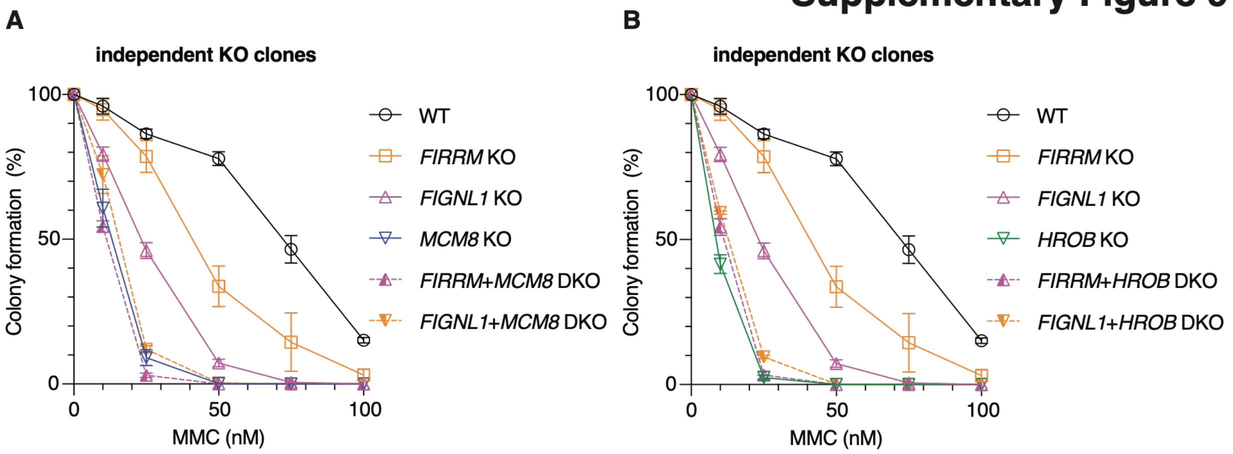


**Supplementary Figure 9. MMC sensitivity of independent *MCM8*, *HROB*, *FIRRM*, and *FIGNL1* KO clones.**

(A, B) Quantification of colony formation by the indicated cells from the three independent experiments (*n* = 3; mean ± SD). The data for WT cells are the same as those shown in Fig. 7B, C, whereas the KO cells were derived from independent clones.
